# Characterization of protein isoform diversity in human umbilical vein endothelial cells (HUVECs) via long-read proteogenomics

**DOI:** 10.1101/2022.05.17.490813

**Authors:** Madison M. Mehlferber, Ben T. Jordan, Erin D. Jeffery, Leon Sheynkman, Jamie Saquing, Bipul R. Acharya, Karen K. Hirschi, Gloria M. Sheynkman

## Abstract

Endothelial cells (ECs) comprise the lumenal lining of all blood vessels and are critical for the functioning of the cardiovascular system. Their phenotypes can be modulated by protein isoforms. To characterize the isoform landscape within ECs, we applied a long read proteogenomics approach to analyze human umbilical vein endothelial cells (HUVECs). Transcripts delineated from PacBio sequencing serve as the basis for a sample-specific protein database used for downstream MS analysis to infer protein isoform expression. We detected 53,836 transcript isoforms from 10,426 genes, with 22,195 of those transcripts being novel. Furthermore, the predominant isoform in HUVECs does not correspond with the accepted “reference isoform” 25% of the time, with vascular pathway-related genes among this group. We found 2,597 protein isoforms supported through unique peptides, with an additional 2,280 isoforms nominated upon incorporation of long-read transcript evidence. We characterized a novel alternative acceptor for endothelial-related gene *CDH5*, suggesting potential changes in its associated signaling pathways. Finally, we identified novel protein isoforms arising from a diversity of splicing mechanisms supported by uniquely mapped novel peptides. Our results represent a high resolution atlas of known and novel isoforms of potential relevance to endothelial phenotypes and function.

**Graphical Abstract:** 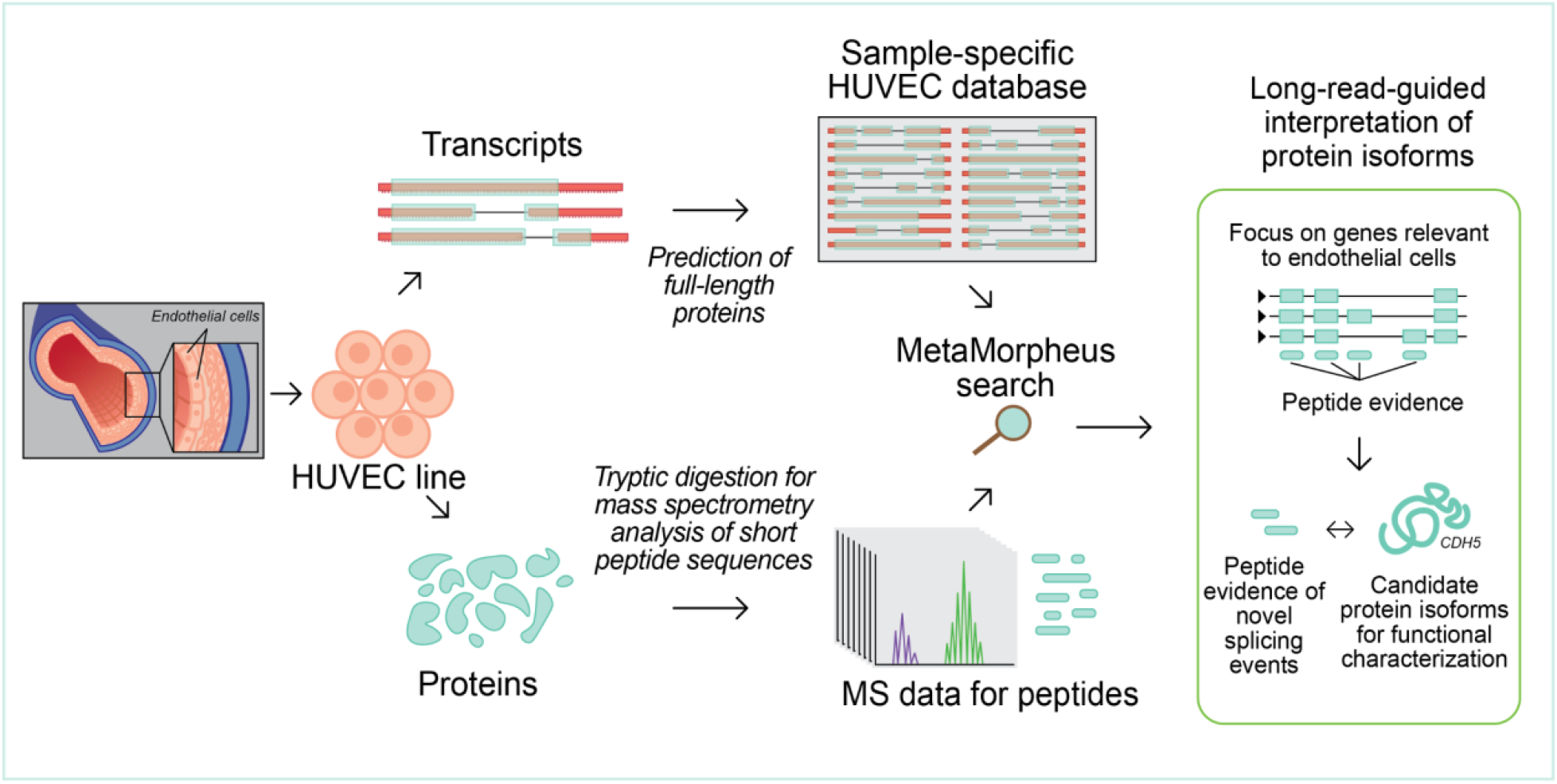

## Introduction

Endothelial cells are critical for the development and maintenance of the cardiovascular system. They form the lining of all blood vessels within the body allowing for functions such as oxygen nutrient delivery, blood pressure regulation, and immune control.^1^ Endothelial dysfunctions can contribute to a host of cardiovascular diseases, such as atherosclerosis, diabetes retinopathy, cancer, and stroke.^2^ Improved understanding of these and related diseases may be attained through molecular characterization of the proteome underlying endothelial cell identity and functionality.^3,4^

Endothelial cells can express functionally distinct protein isoforms through the process of alternative splicing (AS). For example, vascular endothelial growth factor A (*VEGF-A*) exists as two separate isoform families that differentially bind to the extracellular region on VEGFR1 or VEGFR2 leading to proliferation and survival of endothelial cells. One *VEGF-A* isoform family is pro-angiogenic and another is anti-angiogenic.^5,6^ Together these isoforms work in balance to regulate new vessel formation. Globally, across the endothelial cell proteome, many gene functions are modulated by AS.^7–11^ However, despite many high-throughput sequencing datasets collected on endothelial cells^12^, our knowledge of individual protein isoforms that are expressed is incomplete^13^.

In order to characterize the proteome of endothelial cells, human umbilical vein endothelial cells (HUVECs) can serve as a relevant model system, since they are primary cells that can be expanded in culture to generate sufficient material for proteomic analysis.^3,14,15^ A prior study performed by Madugundu and colleagues employed a proteogenomics approach, incorporating RNA-seq and mass-spectrometry (MS)-based proteomics in order to characterize proteomic variation in HUVECs.^16^ By utilizing short-read RNA-seq data, the authors generated a set of custom databases of relevance to isoform detection. Specifically, they generated a database of candidate splice-junction peptides derived from novel exon-to-exon connections (i.e., junctions), as well as a custom database based on inferred reconstruction of full-length transcripts. The study reported both known and novel isoforms, providing further insight into the role of splicing events in HUVECs. However, the proteogenomics approach used relied upon short-read RNA sequencing in the custom database generation, and short reads cannot provide unambiguous knowledge of the *bona fide* full-length isoform (i.e., complete chain of exon/junction connectivities)^17^, which is needed for accurate prediction and detection of full-length protein isoforms.^18^

For improved characterization of protein isoform expression in HUVECs, it would be ideal to obtain full-length transcript information to infer expressed isoforms at the protein level. Fortunately, advances in sequencing technology, such as through the PacBio or Oxford Nanopore long-read sequencing platform, have allowed for detection of full-length transcript isoforms.^19–21^ Capitalizing on these technologies, we previously developed a proteogenomic approach that incorporates long-read RNA sequencing with MS analysis, which we term “long-read proteogenomics”.^22^ Long-read RNA-seq returns information on full-length transcript isoforms^23^, which is bioinformatically translated into full-length protein isoforms predictions.^22,24–26^ These predicted protein isoforms serve as sample-specific, full-length isoform models from which to infer protein expression from MS data.^27^

Here we apply a long-read proteogenomic approach to characterize protein isoforms expressed in HUVECs. We demonstrate the application of PacBio long-read RNA-seq data towards characterization of the full-length transcriptome in HUVECs, which includes detection of unannotated transcript isoforms. A PacBio-derived HUVEC protein database is searched against a sample-matched MS dataset facilitating the characterization of HUVEC-specific isoforms. Finally, we report on the discovery of novel peptides, providing evidence for novel isoforms through a direct mapping of novel peptides to full-length protein isoforms in HUVECs. Overall, we present the first application of a long-read proteogenomics approach as applied to primary endothelial cells. These results nominate candidate isoforms for functional studies of how splicing modulates endothelial cell phenotype and function.

## Experimental Methods

### HUVEC cell culture

Primary Human Umbilical Vein Endothelial Cells (HUVECs) were purchased from Lonza (C2519AS) and used up to passage eight. Early passage HUVECs were cultured in EGM™2- Bulletkit™ medium with growth supplements CC-3156 & CC-4176 purchased from Lonza. At 80% confluency, HUVECs were trypsinized, washed twice with phosphate-buffered saline (PBS), pelleted, and frozen at -80 °C.

### Long-read RNA-seq (PacBio Iso-Seq) library preparation and sequencing run

PacBio (Iso-Seq) data was collected on the HUVEC pellet. HUVEC RNA was analyzed on an Agilent Bioanalyzer to confirm concentration and RNA integrity for downstream analysis. We observed a RIN value of 10. From this RNA, cDNA was synthesized using the NEB Single Cell/Low Input cDNA Synthesis and Amplification Module (New England Biolabs).

Approximately 200 ng of HUVEC cDNA was converted into a SMRTbell library for usage with the Iso-Seq Express Kit SMRT Bell Express Template prep kit 2.0 (Pacific Biosciences). Through this protocol, bead-based size selection occurs in order to remove low mass cDNA (less than 500 kb). Each SMRTBell library was sequenced on the SMRT cell on Sequel II system. A 2 hour extension and 3 hour movie collection time was used for data collection. The “ccs” command from the PacBio SMRTLink suite (SMRTLink version 9) was used to convert raw reads into Circular Consensus (CCS) reads.

### Mass spectrometry-based proteomics sample preparation

Harvested HUVECs, approximately 5 million cells each, were pelleted and frozen at -80 °C. The sample pellet was lysed according to the Filter Aided Sample Preparation (FASP) protocol.^59^ Lysis buffer used in the FASP was changed to 6% SDS, 150 mM DTT, 75 mM Tris-HCl. To the 30 µL pellet of 5 million cells, an aliquot of 60 µL of lysis buffer was added and probe-sonicated to lyse the cells and shear the nucleotide material. Sonication continued for 1-5 minutes until the sample was clear and no longer viscous. The lysate was then incubated at 95 °C for 5 minutes. Protein quantitation was estimated by BCA assay to be approximately 500-600 µg. Quadruplicate aliquots of 20 µL each were subjected to FASP and trypsin digest (1 µg per aliquot) and allowed to incubate at 37 °C overnight. Nanodrop analysis estimated peptide content at 22 µg per trypsin digest (total of 88 µg).

### Offline HPLC Fractionation

The tryptic digests were pooled and dried down to a volume of 40 µL and subjected to offline high pH RP-HPLC fractionation using an Agilent 1200 HPLC. Sample was loaded onto a Thermo Scientific Hypersil Gold C18 column (150 mm x 3 mm x 3 µm C18), equilibrated with 95% solvent A (20 mM NH_4_ formate, pH 10) and 5% solvent B (70% acetonitrile/30% solvent A), and eluted at a flow rate of 400 µL/min, with fractions collected every 1 minute from RT 38-63 min. The following gradient was used: 5% B from 0-30 min, 5-65% B from 30-63 min, 65-100% B from 64-69 min, 100-5% B from 69-70 min, 5% B from 70-73 min. Samples containing peptide, according to UV 214 nm corresponding to the HUVEC pellet were digested with trypsin. Collected fractions 4-20 were selected for LC-MS/MS analysis.

### NanoLC-MS/MS analysis

The resulting peptides were dried to 12 µL and analyzed by nanoLC-MS/MS using a Dionex Ultimate 3000 (Thermo Fisher Scientific, Bremen, Germany) coupled to an Orbitrap Eclipse Tribrid mass spectrometer (Thermo Fisher Scientific, Bremen, Germany). Three microliters of each peptide-containing sample were loaded onto an Acclaim PepMap 100 trap column (300 µm x 5 mm x 5 µm C18) and gradient-eluted from an Acclaim PepMap 100 analytical column (75 µm x 25 cm, 3 µm C18) equilibrated in 96% solvent A (0.1% formic acid in water) and 4% solvent B (80% acetonitrile in 0.1% formic acid). The peptides were eluted at 300 nL/min using the following gradient: 4% B from 0-5 min, 4 to 28% B from 5-210 min, 28-40% B from 210-240 min, 40-95% B from 240-250 min and 95%B from 250-260 min.

The Orbitrap Eclipse was operated in positive ion mode with 1.9 kV at the spray source, RF lens at 30% and data dependent MS/MS acquisition with XCalibur version 4.3.73.11. Positive ion Full MS scans were acquired in the Orbitrap from 375-1500 m/z with 120,000 resolution. Data dependent selection of precursor ions was performed in Cycle Time mode, with 3 seconds in between Master Scans, using an intensity threshold of 2 × 10^4^ ion counts and applying dynamic exclusion (n=1 scans within 30 seconds for an exclusion duration of 60 seconds and +/- 10 ppm mass tolerance). Monoisotopic peak determination was applied and charge states 2-6 were included for HCD scans (quadrupole isolation mode; 1.6 m/z isolation window). The resulting fragments were detected in the Orbitrap at 15,000 resolution with standard AGC target.

### Long-read RNA-seq analysis, MS searching, and proteogenomic analysis conducted using a Nextflow pipeline

The long-read proteogenomics pipeline was implemented with Nextflow, a workflow framework which allows for scalable and reproducible computational analysis. The Nextflow pipeline developed and described previously was used to process HUVEC collected PacBio data, translate the resulting transcripts into the protein database (see *Deriving a HUVEC sample-specific protein isoform database below*), and perform proteomics database searches.^22^ Further information on the workflow including individual modules of the Nextflow pipeline can be found at https://github.com/sheynkman-lab/Long-Read-Proteogenomics.^22^ The GitHub revision (i.e., commit) used in this analysis was https://github.com/sheynkman-lab/Long-Read-Proteogenomics/releases/tag/v1.0.0. All transcriptomic and proteogenomic docker images that are used within the analysis can be found at https://hub.docker.com/ under the repository gsheynkmanlab. The analysis was performed on the University of Virginia High Performance Computing system.

### Long-read RNA-seq (PacBio Iso-Seq) data analysis

The CCS reads obtained from PacBio sequencing were analyzed using the IsoSeq workflow described previously.^22^ Primer removal was done on the 5’ and 3’ end. The 5’ primer consists of an NEB adapter sequence (Sequence: GCAATGAAGTCGCAGGGTTGGG). The 3’ primer consists of the Clontech SMARTer cDNA universal primer (Sequence: GTACTCTGCGTTGATACCACTGCTT). Following processing of the raw reads using the IsoSeq workflow, we derived the number of full-length reads corresponding to each distinct transcript. Full-length read counts per million (CPM) were computed by dividing the number of full-length reads aligning to a transcript isoform by the total number of reads and then multiplying this by a factor of 1,000,000.

### Transcript isoform classification and filtering

SQANTI is a computational tool used for comparison, classification, and quality assessment of the full-length isoforms sequences collected from the long-read platform.^29^ We used SQANTI3 (version 1.3) to annotate the polished transcript isoforms obtained from the Iso-Seq analysis using SQANTI default parameters. *Note*: the default parameters included options to use the genome-derived sequences for the isoform output. As a result, transcriptional variations inclusive of (alternative N-termini, alternative splicing, etc.) but not genetic variations are captured in the HUVEC sample-specific database.

### Generation of a full-length protein isoform database from the long-read RNA-seq

After deriving a high confidence set of full-length transcript isoforms, within the Nextflow pipeline we select the most biologically plausible ORF for each of the Iso-Seq transcripts. Calling the best ORF consists of two steps: finding candidate ORFs (50 nucleotides or longer) using CPAT^60^, and selecting the most plausible ORF based on coding potential, relation of start site to reference start sites, and number of start sites skipped to reach the ORF start site.

To generate the PacBio-derived protein database (HUVEC sample-specific database) employed for downstream MS searching, transcripts were grouped that produced ORFs of the same sequence. The total transcript abundance for each grouping was calculated as the sum of all CPM values for the transcripts comprising that group. Candidate isoforms are further classified based on the protein sequence in relation to the reference protein isoforms, as defined in the ‘sqanti_protein’ and ‘protein_classification’ modules in the Nextflow pipeline. Classifications are based on a variant of nomenclature used within the SQANTI3 software, which we call “SQANTI Protein”.

Additional filtering was performed in order to retain only isoforms that were likely protein coding. Isoforms that did not have a stop codon within the predicted ORF, and could represent truncations, were removed. Isoforms that were either fully mapped to a protein-coding GENCODE reference isoform (“protein full splice match”, pFSM) were retained, as well as isoforms that contained a novel combination of known splice sites or junctions (pNIC). Of the isoforms that contain novel splice sites (pNNC), suspected nonsense mediated decay (NMD) isoforms were removed. Here, NMD suspects were defined as isoforms that contained more than two junctions after the stop codon. Isoforms that were not classified as pFSM, pNIC, or pNNC were removed from consideration. Protein classification details can be found within the ‘protein_classification’ module of the pipeline, while the filtering criteria can be found within the ‘protein_filter’ of the Nextflow pipeline.

A hybrid database was developed that incorporated isoforms from PacBio if the gene resided in the high confidence region, defined as where the aggregated transcriptomic gene abundance contained at least three CPM and the average reference exonic gene length was between 1 and 4 kilobases (kbp). If a gene did not meet this criteria the reference isoforms were substituted in place of the long-read isoforms. If a gene was not found within the long-read transcriptomic data the reference protein isoforms were also appended into the hybrid database. A detailed description of reasoning behind creation of a hybrid database has been described previously.^22^

### GENCODE and UniProt reference protein database

The GENCODE protein database used in this study was created by downloading the coding translation FASTA and grouping entries with the same protein sequence for each gene (‘make_gencode_database’ module in the Nextflow). For the many cases where one or more GENCODE transcripts from the same gene lead to the same protein sequence, the transcripts were grouped and given a defining protein accession as the first alphanumeric GENCODE protein accession, by the transcript name (GAPDH-201).

The UniProt database used was the reviewed human database with isoforms, downloaded November 1st, 2020. The database contains 42,358 protein isoform entries from 20,292 genes.

### MS database search

Standard proteomic analysis of acquired mass spectra files were performed using the free and open source search software program MetaMorpheus.^43^ A custom branch and Docker image were made as part of the nextflow piplemen based GitHub: https://github.com/smith-chem-wisc/MetaMorpheus/tree/LongReadProteogenomics, Docker: https://hub.docker.com/r/smithchemwisc/metamorpheus/tags?page=1&ordering=last_updated tag: lrproteogenomics) based on MetaMorpheus version 0.0.316. Analysis of the collected spectra files performed either using the HUVEC sample-specific database (HUVEC-derived PacBio reads + GENCODE entries; “HUVEC sample-specific database”) (71,511 of entries from 19,982 genes) in which the subset of PacBio derived entries are 26,675 protein isoforms from 7,283 genes. The GENCODE human database (version 35; 87,729 protein entries from 19,982 genes), or the UniProt reviewed human database with isoforms (downloaded July 8th 2021; 42,380 protein entries from 20,292 genes). All searches were conducted with a contaminants database, included in MetaMorpheus, which contains 264 common contaminant proteins frequently found in MS samples.

All RAW spectra files were first converted to MzML format with MSConvert prior to analysis with MetaMorpheus (see ‘mass_spec_raw_convert’ module in the Nextflow pipeline). For the MetaMorpheus MS search, the settings used for all search tasks can be found in Supplementary Table 5. MetaMorpheus produces peptide spectral match (PSM), peptide and protein group result files, which we analyzed in downstream custom modules. All peptide and protein results reported employ a 1% False Discovery Rate (FDR) threshold after target-decoy searching.^42^

### Criteria for Novel Peptide Identification

Stringent filtering criteria and manual validation was used, as described previously^22,61^ to ensure that the spectrum does in fact represent the novel peptide sequence. Spectra corresponding to the scan number of the identified novel peptide sequence were derived from MetaDraw and manually inserted into an Excel file which were then manually evaluated. In addition to previously^22^ described criteria for novel peptide annotation, additionally, we allowed for cases where the C13 isotope for a novel peptide.

### Data analysis and plot generation

All downstream data analyses were performed through custom Python scripts. Data analysis scripts used for generation of figures, plots, and statistics may be found in the following GitHub repository: https://github.com/sheynkman-lab/Huvec-Proteogenomic-Analysis

### Availability of data and materials

Raw long-read RNA-seq data collected on the PacBio platform are available from the Sequence Read Archive (PRJNA832812, corresponding to accession SRR18959149). Data generated by math spectrometry are available through MassIVE, the Mass Spectrometry Interactive Virtual Environment (MSV000089326, username: MSV000089326_reviewer, password: sheynkmanhuvec). The output of the data analysis including the long-read proteogenomics Nextflow workflow results generated using the mass spectrometry and long-read RNA-sequencing data as well as the post pipeline analysis results are available on Zenodo (10.5281/zenodo.648486).

The open-source software produced in the making of this work is freely available under the MIT license found in the GitHub repository (https://github.com/sheynkman-lab/Long-Read-Proteogenomics). A wiki was created (https://github.com/sheynkman-lab/Long-Read-Proteogenomics/wiki) describing each of the pipeline processes.

Code used to generate the main figures and tables in this manuscript can be found in the GitHub repository (https://github.com/sheynkman-lab/Huvec-Proteogenomic-Analysis).

## Results

### Long-read proteogenomics to characterize isoforms in endothelial cells

In order to characterize the isoforms expressed in an endothelial cell population, we subjected HUVECs to “long-read proteogenomics” where samples undergo long-read RNA-sequencing and mass-spectrometry analysis in parallel, which is followed by integrative analysis of the matched datasets.^22^ The full-length transcripts—obtained from long-read RNA-seq—are converted to a predicted protein database, serving as candidate isoforms for proteomic detection (Figure 1). As a first step in our method, PacBio RNA sequencing is performed to characterize the HUVEC transcriptome.

**Figure 1.**
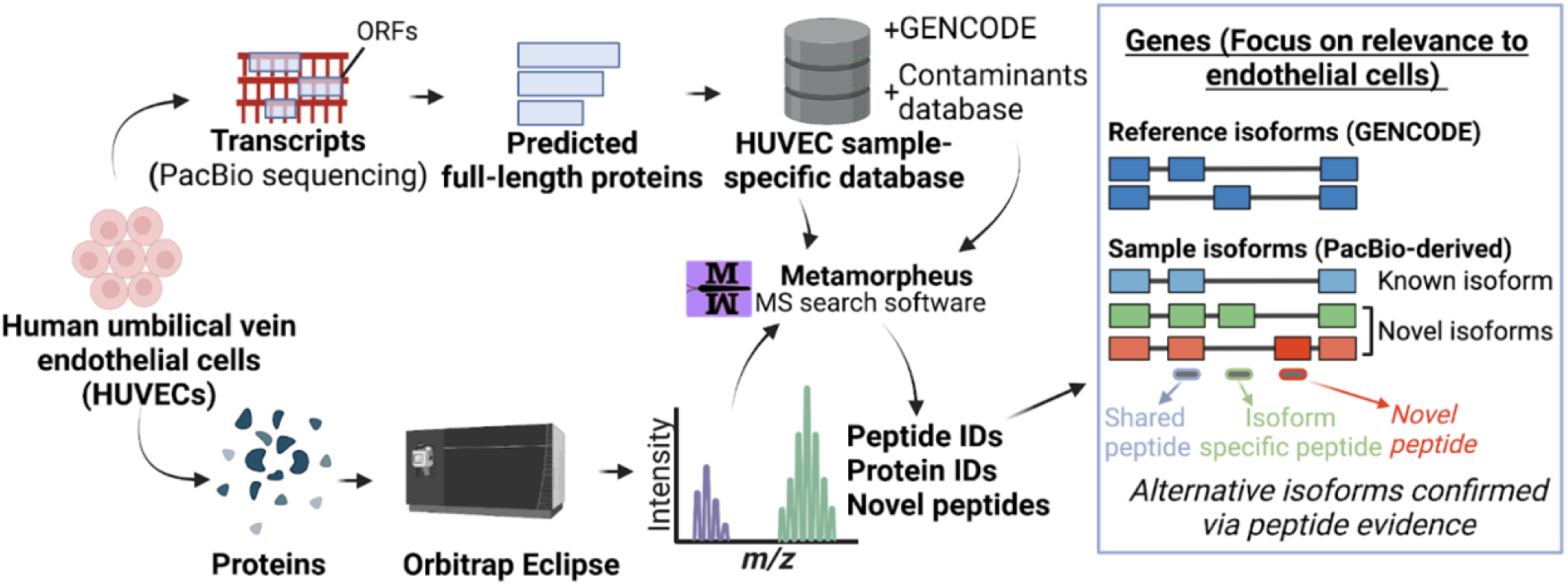
Characterization of isoform diversity in HUVECs through integration of long-read RNA-seq with mass spectrometry data (“long read proteogenomics”). Transcripts are converted into a protein isoform database based on predicted open reading frames (ORFs) and the resulting database is searched against a sample-matched bottom-up mass spectrometry (MS) dataset. The peptide identifications can be used to support the expression of isoform candidates related to endothelial pathways.

### Long-read RNA-seq of HUVECs reveals widespread and novel isoform diversity

Long-read RNA-seq data was collected on the PacBio sequencing platform using the “Iso-Seq” method^28^, generating 3,608,972 long-reads (i.e., circular consensus reads). These reads were processed by Iso-Seq3^28^ to generate the set of distinct transcript isoforms and their respective abundances (Figure 2A).^22^

**Figure 2.**
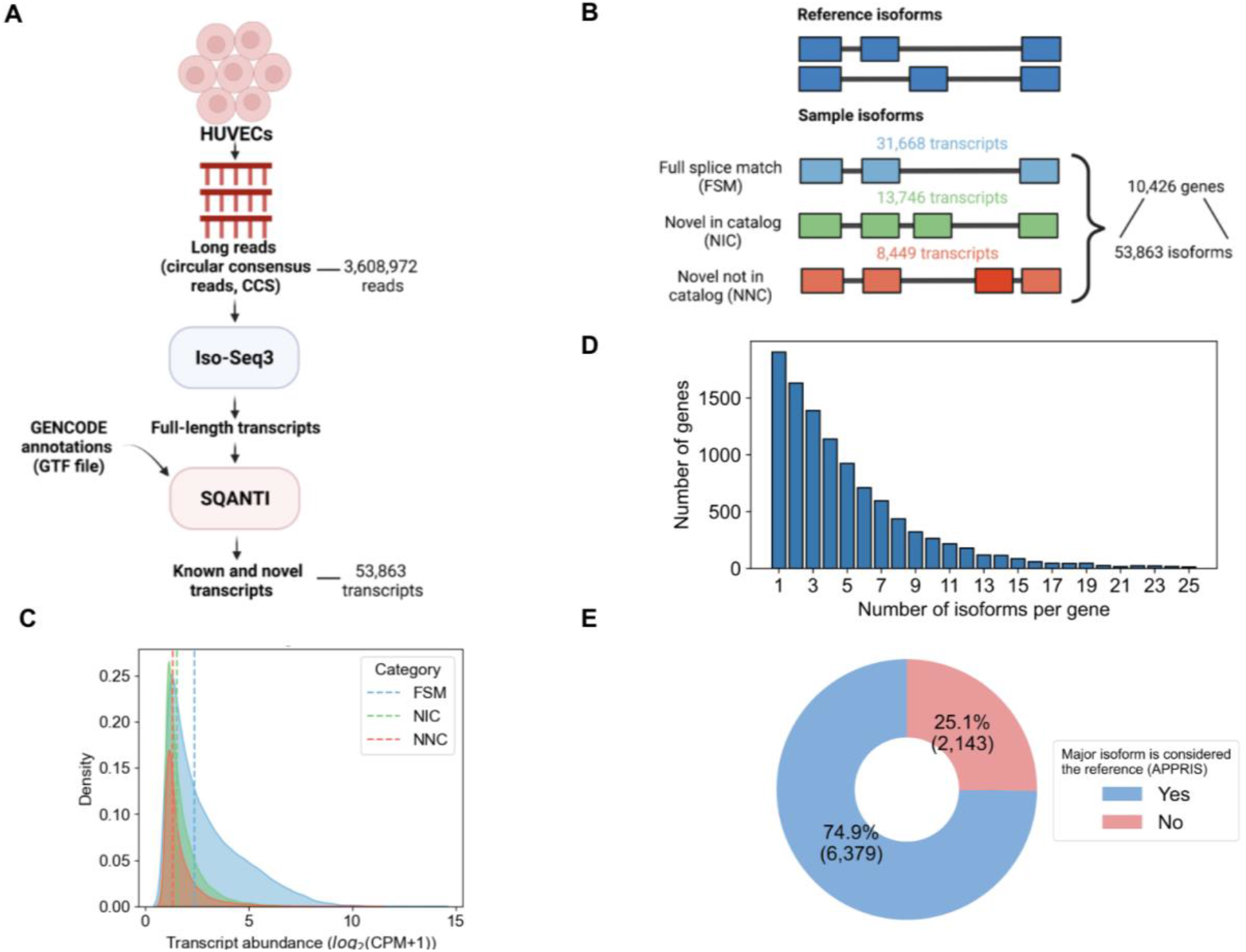
Characterization of transcript isoform diversity in HUVECs via long-read RNA-Seq. (A) Schematic of the long-read RNA-seq analysis pipeline. (B) Transcripts and genes identified from PacBio long-read RNA-seq. The number of known (blue) and novel isoforms (green and orange) are shown. (C) Transcript abundance distribution for known (FSM) versus novel transcripts (NIC, NNC), with dashed lines representing median abundance values in full-length read counts per million (CPM) for each category (FSM= 2.36, NIC=1.5, NIC=1.3). (D) Distribution of the number of genes expressing multiple isoforms. (E) Fraction of genes in which the most abundantly expressed isoform (“major isoform”) differs from the reference isoform (APPRIS principal isoform).

PacBio-derived transcripts were compared to reference transcripts (GENCODE v35) and their novelty status was defined using SQANTI3 (Figure 2B).^29^ The UniProt database lacks a complete mapping of protein isoforms to the reference genome, and therefore we could not compare transcripts to UniProt directly, although future efforts may resolve this limitation.^30–32^ Based on a comparison to GENCODE models, we identified 53,863 transcripts from 10,426 protein coding genes, inclusive of all transcripts with a minimum abundance of one full-length read count per million (CPM). The average length of transcripts is 2,846 kilobase pairs (kbp) (Supplementary Information Figure S1A). Among the 53,863 transcripts isoforms identified in the HUVEC sample, 31,668 (59%) matched exactly to a transcript isoform in GENCODE, the match being based on splice junction connectivity (“full splice matches”, “FSM”, Figure 2B). The remaining 22,195 (41%) isoforms were unannotated, or novel, in terms of the observed ordering of splice junctions along the length of the transcript (Figure 2B). Of the unannotated isoforms identified, 13,746 (62%) contained novel combinations of known splice junctions (“novel in catalog”, “NIC”), and the remaining 8,449 (38%) isoforms contained entirely new exon splice boundaries, in which the acceptor or donor site is not represented in GENCODE (“novel not in catalog”, “NNC”, Figure 2B). The overall abundance distribution for identified transcripts was wide ranging (see Supporting Information Figure S1B). As expected, on average, the novel transcripts exhibit lower abundance than known transcripts (Figure 2C).^33,34^ The FSM transcripts displayed a median abundance of 2.4 CPM, while the NIC and NNC transcripts displayed a median of 1.5 and 1.3 CPM, respectively. This data illustrates that novel transcripts tend to exhibit lower abundances than known transcripts. While these trends represent average expression differences, particular novel transcripts can exhibit high abundances within HUVECs.

Using the full-length transcriptomics dataset, we next determined the number of protein-coding genes that returned evidence for expression of multiple isoforms. We found that 82% (8,522 genes of the 10,426 genes represented) of detected genes expressed multiple transcript isoforms (Figure 2D). To focus on genes involved in endothelial pathways that may be co-expressing multiple isoforms, we manually curated the literature to compile a list of genes that are involved in vascular pathways related to early endothelial differentiation and development or hemogenic specification (see Supporting Information Table S1).^35,36^ We then determined which endothelial genes are expressing multiple isoforms in our HUVEC sample. To have increased confidence in isoform expression of such genes, we filtered for genes which contain two or more isoforms with each isoform having an abundance of at least three CPM. We identified multiple co-expressing isoforms for *CD34, CELF1, FLT1, NRP1* and *SRSF5* (Table 1, with annotations from GOrilla^37^). Notably, we discovered novel isoforms for neuropilin or *NRP1*, a gene that expresses proteins involved in regulating vascular morphogenic pathways through its binding interactions with VEGF-A.^38^ Interestingly, this pathway is functionally modulated by different isoforms of VEGF-A.^6,39^

**Table 1.**
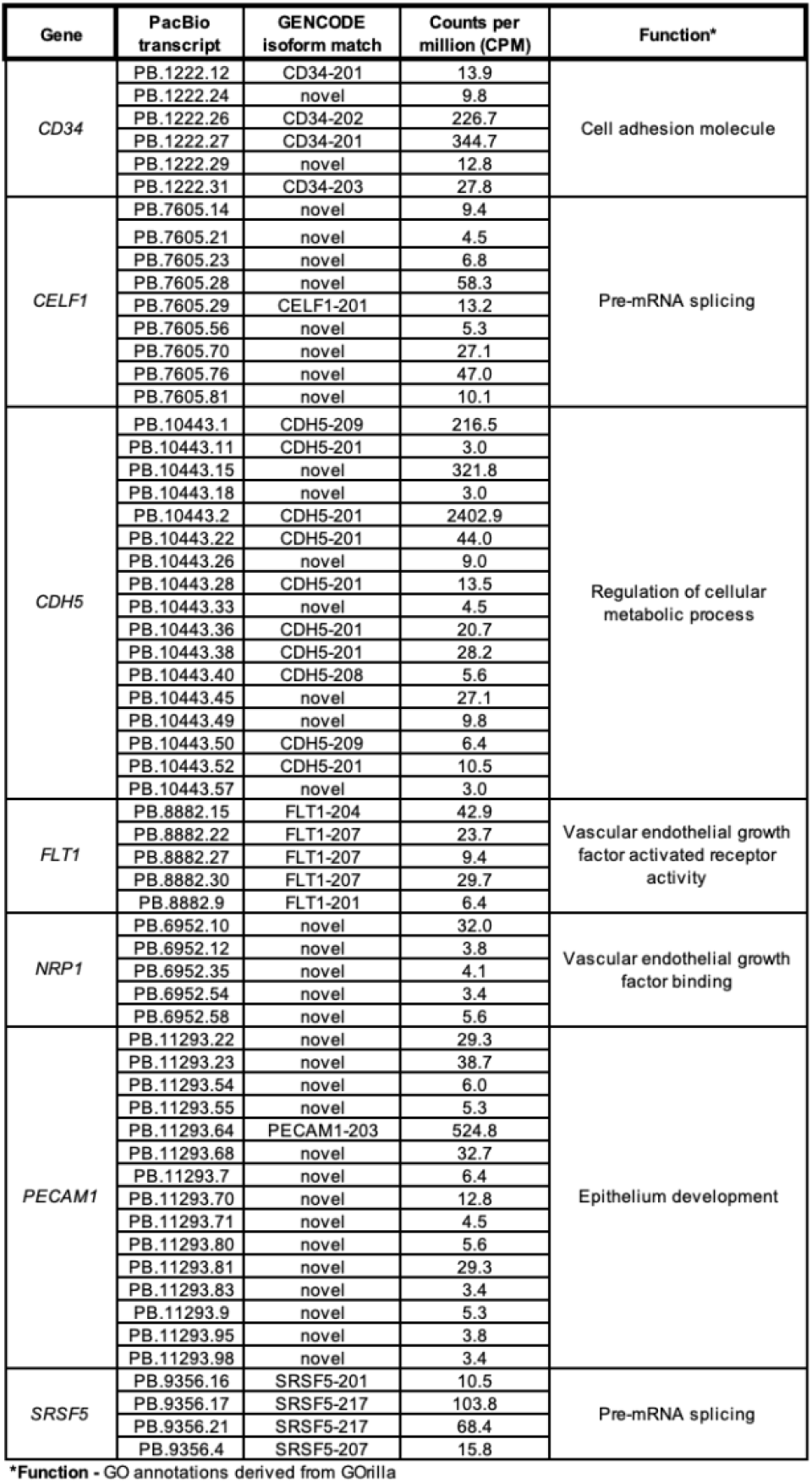
Endothelial-relevant genes expressing multiple transcript isoforms in HUVECs.

Given the prevalence of genes that co-express multiple isoforms in HUVECs, we next asked to what extent the identity of the most highly expressed isoform, i.e., the major isoform, matches what is defined as the “reference isoform” for a gene. To define a gene’s “reference isoform”, we used the APPRIS database which reports a principal isoform to be most representative for a gene.^40^ The APPRIS principal isoform concept is related to the concept of a UniProt “canonical” protein, though underlying assumptions differ.^31,40^ For the genes expressing multiple isoforms, we classified their corresponding isoforms as either major, i.e., the most abundant isoform based on relative expression levels of all isoforms for a gene, or minor. There were 1,904 genes only expressing one isoform and therefore were excluded from analysis. We identified 8,522 transcripts as the major isoform and 43,437 as minor isoforms. We found, as expected, that on average the major isoforms are more highly expressed than minor isoforms (Supplementary Information Figure S1C-D). Surprisingly, we found that for 25% (2,143 isoforms) of genes, the major isoforms in our HUVEC sample do not coincide with the APPRIS principal isoform (Figure 2E). Within this population of major isoforms, we found six genes involved in endothelial pathways, *CELF1, FLT1, GATA2, NR2F2, NRP1, NRP2* and *SRSF6* (see Supporting Information Table S2). These results illustrate that the major isoform expressed in a given sample may not always correspond to the generic “reference” isoform for a gene, which can be explained by the fact that isoforms exhibit cell or tissue-specific expression patterns.^41^

Collectively, the HUVEC transcriptomic results demonstrate the use of long-read RNA-seq to characterize sample-specific variation in isoform identity and abundance.

### Deriving a HUVEC sample-specific protein isoform database

The vast transcriptome diversity of HUVECs likely translates in some part to a diversity of protein isoforms. To explore this question, we translated the HUVEC transcript isoform sequences *in silico* into open-reading frames (ORFs) and compiled the predicted sequences into a HUVEC sample-specific protein isoform database for MS searching, as previously described (Supplementary Information Figure S2A).^22^ To classify the relationships between the predicted proteins to that of annotated protein isoforms in GENCODE^30^, we used the classification scheme we previously developed, SQANTI Protein.^22^ SQANTI Protein automatically categorizes known and novel protein isoforms. The categories include “protein full-splice match” (pFSM), “protein novel in catalog” (pNIC), and “protein novel not in catalog” (pNNC) (Supplementary Information Figure S2B). We found that 16,296 predicted proteins exactly matched protein isoforms in the GENCODE reference (pFSMs), while 24,896 predicted protein isoforms were novel (Supplementary Information Figure S2C). Among those novel isoforms, 5,855 had novel combinations of known protein sequence elements such as the N-terminus, the C-terminus or the splicing pattern (pNICs). The other 19,041 protein isoforms had one or more entirely novel elements, such as a novel N-terminus or an unannotated exon (pNNC).

Among the candidate protein isoforms, we first filtered out protein isoforms that may have resulted from transcripts from incomplete reads or poor-quality transcripts (see *Protein database generation* in Methods; 11,876 filtered out). The remaining 34,531 predicted protein isoforms (comprising 16,296 pFSMs, 5,855 pNICs, and 12,389 pNNCs) from 10,912 genes were compiled to create a preliminary HUVEC protein database (Figure 3A). These genes and their associated isoforms represent candidates for inclusion in the final database. For the final database, we decided to only include isoforms from genes for which we could ensure a complete sampling of the transcripts, and thus the predicted proteins. Therefore, we created a hybrid database in which we defined a core set of genes for which the transcript detection, and thus predicted proteins, is likely complete based on the long-read data collected. The core set of genes included in the hybrid database have a minimum abundance of three CPM and a moderate transcript length (1-4 kbp average GENCODE-annotated transcript length). For all other genes, the hybrid database is populated with all GENCODE protein isoform entries. The hybrid structure of the final database ensures comprehensiveness of the protein models, with the protein completeness assumption of target-decoy searching satisfied so as to avoid issues of an off-target peptide match.^42^

**Figure 3.**
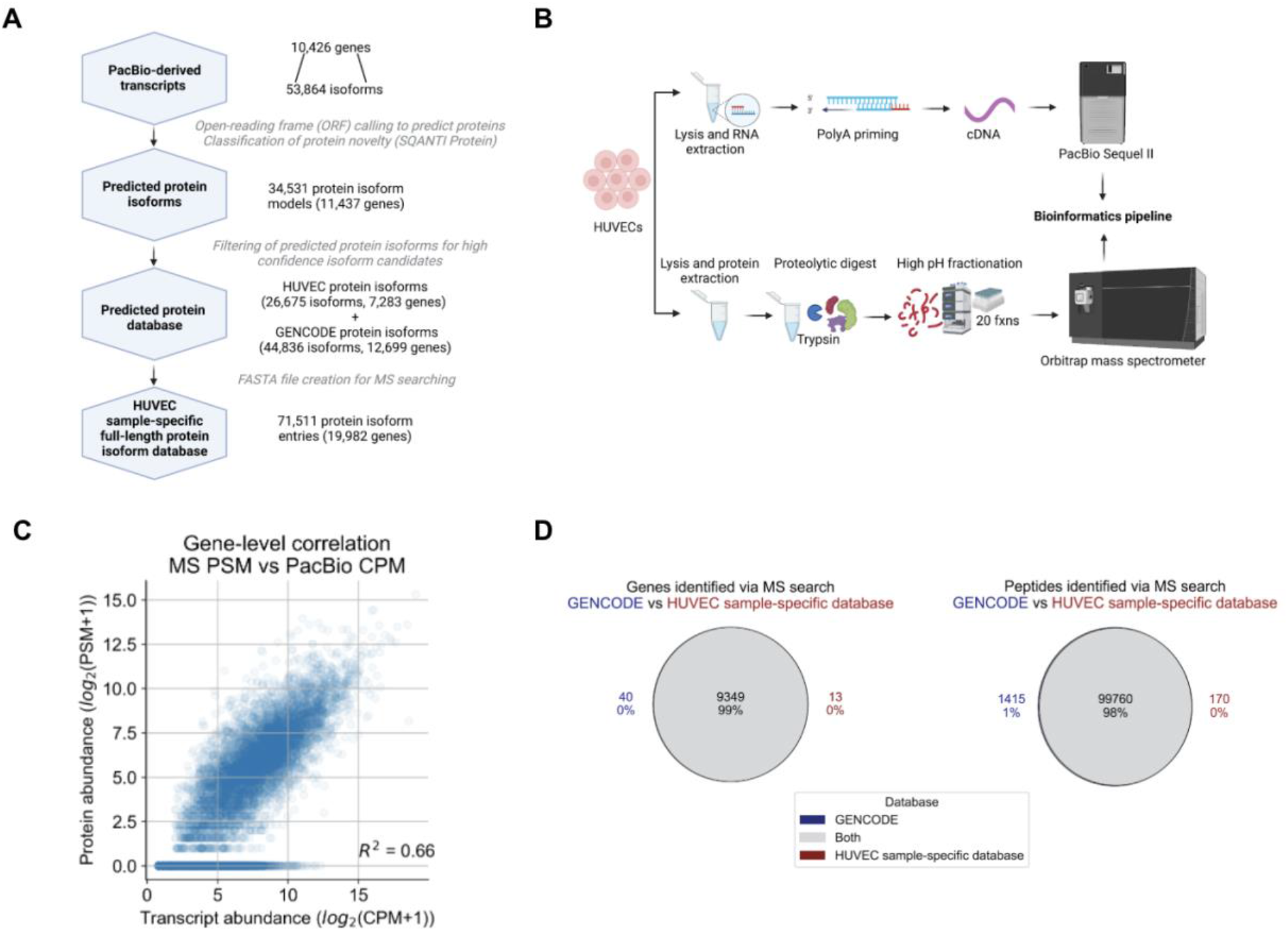
Proteomic analysis of HUVECs using a customized long-read-derived protein isoform database. (A) Steps involved in the generation of a HUVEC sample-specific database. (B) Parallel long-read RNA-seq and MS proteomic data collection from HUVECs. (C) Correlation between estimated RNA and protein expression levels. PSM, peptide spectral match; CPM, full-length read counts per million. (D) Comparison of proteomic search results between the reference and HUVEC sample-specific database.

As described, the final HUVEC sample-specific database for proteomic analysis includes a mixture of custom PacBio-derived proteins as well as annotated GENCODE proteins (Table 2). A detailed listing of steps to convert the transcriptome data to a protein database may be found in Supplementary Information Table S3.

**Table 2.** Composition of the HUVEC sample-specific database.

| PacBio-derived HUVEC sample-specific database |  |  |
| --- | --- | --- |
| Source | Genes | Protein entries |
| GENCODE | 12,699 | 44,836 |
| PacBio-derived (HUVECs) | 7,283 | 26,675 |
| Contaminants* | - | 264 |
| Total | 19,982 | 71,511 |

\*Derived from the MetaMorpheus software version 0.0.316

### Collection of a deep-coverage MS dataset for HUVECs

In order to characterize protein isoforms in HUVECs, we generated and analyzed a deep-coverage MS dataset collected on the same HUVEC pellets that were used for long-read RNA sequencing (Figure 3B). HUVECs were lysed and processed using the filter aided sample preparation (FASP) protocol, in which protein was digested with trypsin to generate a mixture of tryptic peptides. The tryptic digest was subjected to off-line fractionation on an analytical scale high-pH reverse-phase liquid chromatography instrument, and 20 fractions were collected (Supplementary Information Figure S3). These fractions were then analyzed via liquid chromatography LC-MS/MS (Orbitrap Eclipse) in data-dependent acquisition (DDA) mode, generating 3,772,771 MS2 fragmentation spectra. Acquired spectra were searched using the MetaMorpheus^43^ software to obtain peptide and protein identifications passing a 1% false-discovery-rate (FDR). Parameters for the MS search can be found in Supplementary Information Table S4.

### The HUVEC-specific protein database returns near-complete coverage of detectable peptides from a reference search

To use PacBio-derived transcripts as the basis for deriving a protein database for MS searching, a key assumption is that abundances of transcripts are at least moderately correlated with protein abundance. In the past, moderate RNA-protein correlations have been observed using short-read RNA-seq or microarray datasets to quantify transcript abundance.^44^ Here, we examined the correlation of the transcript abundance that is computed from the long-read RNA-seq data (sum total transcript abundance for a gene, in units of CPM) to the estimated protein abundance (sum total peptide counts passing a 1% FDR, in units of number of peptide spectral matches or PSMs). We observed a moderate correlation with an R-square of 0.66 (Figure 3C), providing support that the PacBio-based transcript abundances should serve as a reasonable proxy for protein presence, although that may not always be the case for a particular gene.

To assess the general protein sequence content of the HUVEC sample-specific database (not resolved to individual isoforms), we assessed recovery of annotated peptides and genes. The MS data was searched against the GENCODE and UniProt databases to define the set of annotated peptides and genes detectable in the HUVEC sample, and then the same data was searched against the HUVEC sample-specific database. We found that the HUVEC sample-specific database search returned 98% of the peptide and 99% of the gene identifications that were identified when using the GENCODE database for searching (Figure 3D). The extent of overlap between peptides and genes was similar for the UniProt search results (Supplementary Information Figure S4). Overall, these results indicate that the HUVEC sample-specific database, which was derived *de novo* from long reads, is able to capture a majority of the detectable gene and peptide populations likely expressed in HUVECs. Confirmation of the large overlap of peptide populations identified by the sample-specific database is ultimately useful since it is the underlying populations of peptides identified that are the basis for protein isoform characterization.

### Characterization of HUVEC protein isoforms based on available peptide evidence

We have shown that nearly all reference annotated peptides that are detectable are represented in the HUVEC sample-specific database. With the goal of characterizing isoform expression in endothelial cells, we next evaluated the evidence for the presence of isoforms, in terms of the patterns of their underlying peptide identifications. Due to the complexities and potential ambiguities of protein inference^27^, we elected to examine the peptide evidence directly.

We defined three scenarios of isoform detection precision, based on how the set of identified peptides map to isoforms of a gene. The first scenario is when all isoforms of a gene contain only shared peptides, in which the presence of any isoform cannot be definitively confirmed (Figure 4A, “Protein isoforms correspond to shared peptides”). Among the 10,444 genes with any peptide evidence, we found that 5,993 genes (57%) are cases in which no isoforms for the gene can be specifically confirmed as expressed because all mapped peptides are shared among two or more isoforms. Of these genes evidenced only by shared peptides, 3,436 are genes containing PacBio-derived protein isoforms in the hybrid database.

**Figure 4.**
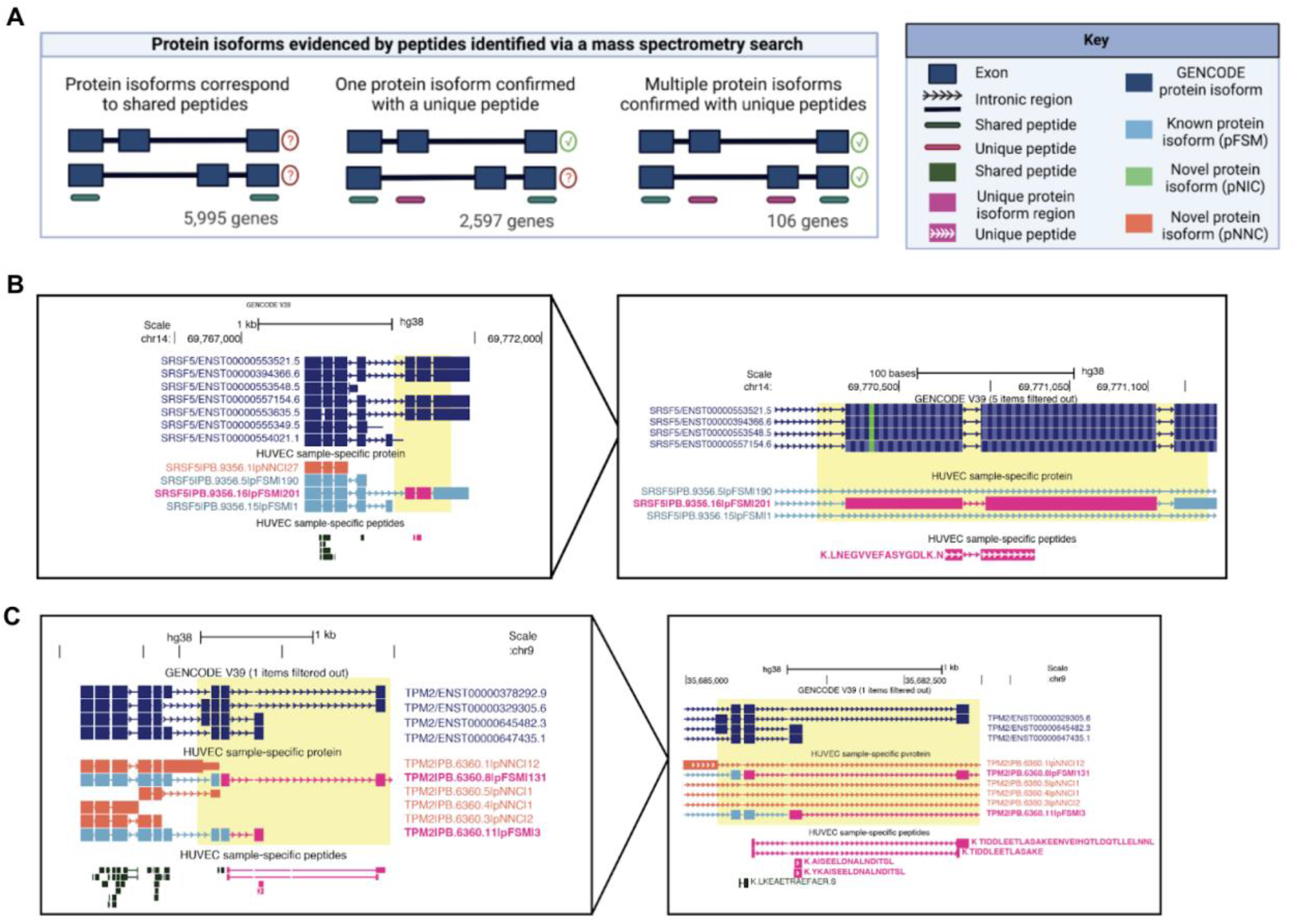
Protein isoforms analyzed based on peptides identified via mass-spectrometry (MS). (A) Scenarios of differing protein isoform detection precision when evidenced by peptides identified from MS. Only genes with multiple protein isoforms in the database are included, and 1,904 genes that express only one isoform were excluded. (B) A protein isoform confirmed with a uniquely mapping peptide LNE, for *SRSF5*, a splice factor that regulates transcripts of *VEGF-A*. (C) Two protein isoforms of *TPM2* are confirmed with uniquely mapping peptides TID, AIS, and YKA. In B and C PacBio-derived protein isoform label follows this format: <Gene>|<PB accession>|<SQANTI Protein class>|<CPM>.

In all other scenarios, there is evidence for the existence of a specific protein isoform because one or more isoforms contain a uniquely mapping peptide. Indeed, the second scenario is when an isoform-specific peptide is identified (Figure 4A, “One protein isoform confirmed with a unique peptide”). We found 4,451 (42%) genes for which we have unambiguously identified at least one isoform for a gene. For 1,748 (17%) of genes, only a single isoform was listed in the database, thus, all peptides would be expected to be uniquely mapped. For the remaining 2,703 (26%) of genes with multiple isoforms annotated, 2,597 (25%) of genes have a single isoform with unique peptide evidence. For example, we found a single isoform supported by a uniquely mapping peptide (Sequence: LNEGVVEFASYGDLK) for Serine and Arginine Rich Splicing Factor 5 (*SRSF5*), which is involved in the splicing of *VEGF-A* pre-mRNA (Figure 4B). Notably, this peptide is shared among two isoforms in the GENCODE database, meaning that the reference database search results cannot pinpoint the source isoform for this peptide.

Of particular interest is a third scenario in which we found evidence for co-expression of two or more isoforms, each supported by a uniquely mapping peptide. In such cases, a natural question is the nature of the functional relationship between the two isoforms and their biological role in endothelial cells. We found 106 (1%) genes with evidence of two or more co -expressing isoforms (Figure 4A, “Multiple protein isoforms confirmed with unique peptides”). For example, we found two isoforms for Tropomyosin 2 (*TPM2*), each supported by a unique peptide (Figure 4C). Notably, there were nine genes in which three or more isoforms each had unique peptide evidence. Interestingly, there was an unusually large number of seven protein isoforms detected from the gene Plectin (*PLEC*), which exhibits a series of alternative N-termini due to differential 5’ transcription (Supporting Information Figure S5). A list of all protein isoforms supported by peptide evidence can be found in Supplementary Information Table S5.

Collectively, these results highlight that while some isoforms may be readily identified from peptide evidence alone, overall, the standard bottom-up MS approach alone does not reach the coverage needed to directly characterize all isoforms predicted from the transcriptome, as observed previously.^45,46^

### Increased support for protein isoform presence in HUVECs through incorporation of underlying transcript evidence from long-read RNA-seq

Despite the use of a sample-specific protein isoform database for MS analysis, a large population of predicted isoforms are only supported by shared peptides (Figure 4A). Because shared peptides map ambiguously to multiple isoforms, they cannot directly confirm expression of any particular isoform in the sample. However, the evidence for a particular protein isoform could be strengthened by considering the underlying transcript abundance levels provided by the sample-matched long-read RNA-seq data, a concept we previously introduced, as well as described for short-read RNA-seq data.^22,47,48^ We reasoned that transcript abundance could be used as an additional source of evidence in the isoform discovery process, given there is a moderate correlation between RNA and protein abundance (Figure 3C).

To explore how long-read RNA-seq data can nominate particular protein isoforms, we first focused on scenarios for which all predicted isoforms for a gene are supported only by shared peptide support. Among such ambiguous protein isoform sets, we reasoned there is higher likelihood for expression of protein isoforms for which the associated transcript abundance is moderately high (e.g., 25 CPM or higher, Figure 5A). As described in the previous section, 5,993 genes had only shared peptide evidence. Among those genes, 3,436 (57%) contained PacBio-derived isoforms, which have associated transcript abundance information.

**Figure 5.**
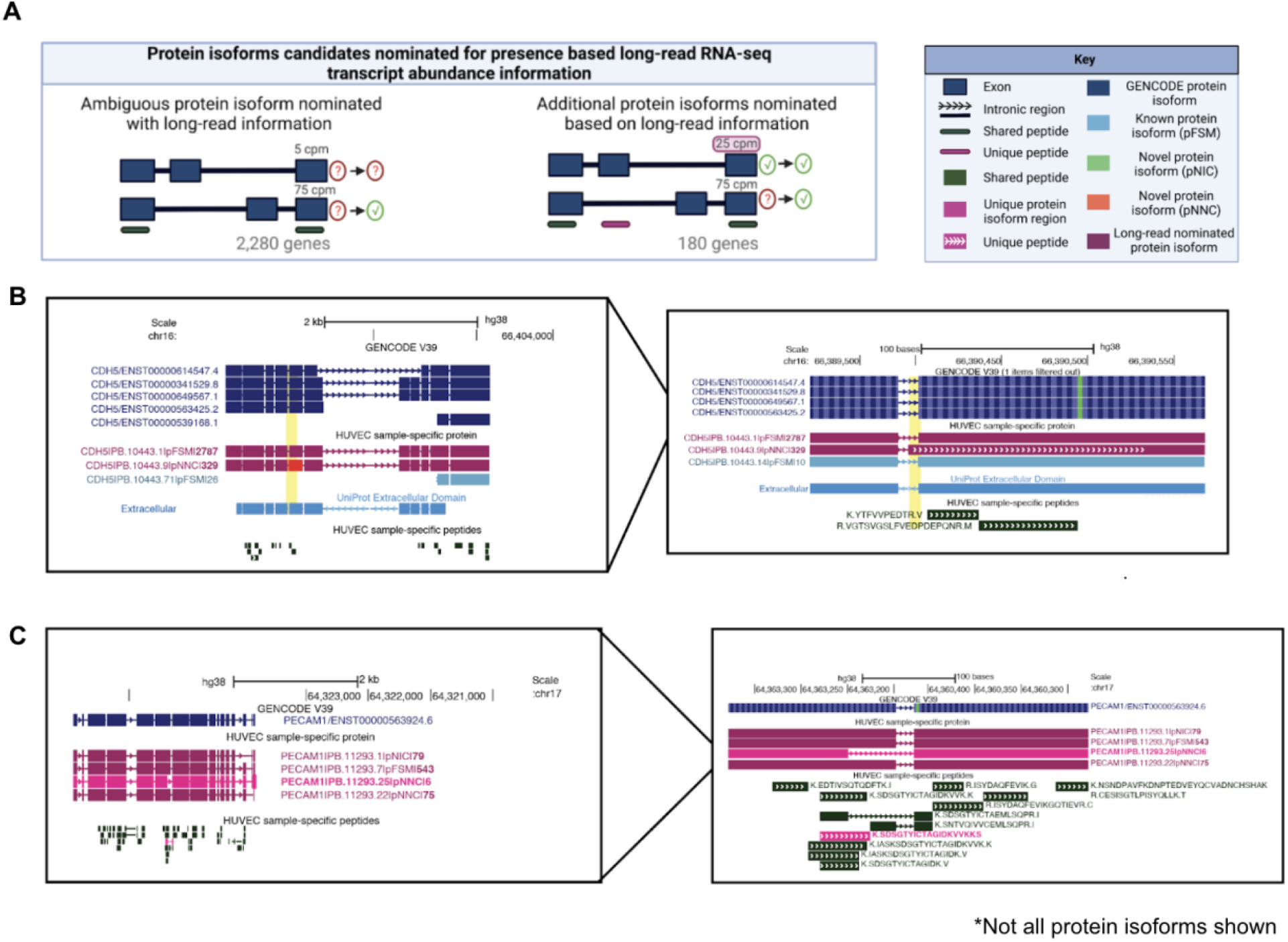
Nomination of protein isoforms when incorporating long-read data. (A) Scenarios of protein isoform candidates nominated for expression when transcript abundance from the long-read RNA-seq information is incorporated. (B) *CDH5* gene, involved in endothelial pathways demonstrating a scenario of ambiguous protein isoforms identified only by shared peptides, but incorporation of long-read RNA-seq data suggests the expression of three moderately expressed protein isoforms (PB.10443.1, PB.10443.9 and PB.10443.71). (C) *PECAM1* gene, involved in endothelial pathways demonstrating an example where one protein isoform is identified via a unique peptide (PB.1123.25), SDS, while the remaining protein isoforms are supported by shared peptides. Abundance information from long-read RNA-seq suggest expression of (PB.11293.1 and PB.11293.7). In B and C, PacBio-derived protein isoform label follows this format: <Gene>|<PB accession>|<SQANTI Protein class>|<CPM>. For B and C, low abundance protein isoforms (<25 CPM) are not shown.

We found that 2,280 (38%) out of the 3,436 genes contain at least one isoform with a moderately high transcript abundance of 25 CPM or higher (Figure 5A, “Ambiguous protein isoform nominated with long-read information”). Interestingly, we found 247 (4%) genes in which there is potential co-expression of at least two protein isoforms in HUVECs. For example, we found that *CDH5*, otherwise known as VE-Cadherin^49^, potentially expresses multiple protein isoforms. One isoform is highly expressed (PB.10443.1; 2,787 CPM) and matches a protein isoform in GENCODE and UniProt (GENCODE isoform CDH5-201, UniProt accession P33151). However, another isoform is robustly expressed (PB.10443.9, 326 CPM) and, interestingly, is a novel isoform because of alternative usage of a novel splice acceptor (Figure 5B). This splicing event leads to an isoform of *CDH5* that gains nine amino acids in the extracellular domain, the region of the protein responsible for mediating interactions with other cadherins to regulate endothelial adhesion properties. This example highlights that while *CDH5* isoforms were only supported by shared peptides, the inclusion of the transcript abundance information as provided by the matched long-read data provides higher weights on the existence of at least two isoforms.

To further explore how long-read RNA-seq data can provide additional evidence for expression of protein isoforms, we focused on scenarios in which there is clear evidence for one isoform based on unique peptide evidence, but another isoform of the same gene is supported by only shared peptides (Figure 5A, “Additional protein isoforms nominated based on long-read information”). We found 180 genes (3%) for which the existence of the alternative protein isoform is supported by long-read evidence (i.e., 25 CPM or higher transcript abundance). Interestingly, we found several protein isoforms of a key endothelial cell surface marker, the platelet endothelial cell adhesion molecule, *PECAM1* (also known as *CD31*).^50^ We found a unique peptide identified for *PECAM1* (PB.11293.25, Sequence: SDSGTYICTAEMLSQPR), but the remainder of peptides identified for PECAM1 are shared across multiple *PECAM1* isoforms, leaving open uncertainty about the expression of other *PECAM1* isoforms beyond PB.11293.25. From the transcript abundance information, we nominated three additional isoforms accompanied by strong long-read support for *PECAM1* (PB 11293.22, 75 CPM: PB 11293.1, 79 CPM; PB 11293.7, 543 CPM; Figure 5C). *PECAM1* produces a transmembrane protein with an extracellular domain, transmembrane-spanning domain, and a C-terminal cytoplasmic domain that likely interacts with intracellular signaling proteins in endothelial cells.^50–52^ Strikingly, the differential exon usage observed for these three isoforms are located exclusively in the C-terminal domain, suggesting potential changes to interactions with intracellular signaling molecules. Further details on candidates identified via long-read abundance information can be found in Supplementary Information Table S5.

Collectively, these case studies highlight how inclusion of transcript abundance information could nominate protein isoforms which were unable to be directly confirmed as expressed based solely on MS peptide evidence. Note that this approach does not provide any information on the absence of protein isoforms with lower transcript abundance, but, rather, is supplying additional lines of evidence to nominate protein isoforms that may have higher likelihood of expression and represent candidates for functional study. Such isoforms are attractive candidates for further MS validation and subsequent functional analysis.

### Novel protein isoform discovery enabled through the HUVEC sample-specific database

We have shown that utilization of a HUVEC sample-specific protein database, with the accompanying transcript abundance values, can lead to inference of novel protein isoform presence. A more direct way to confirm the presence of a novel protein isoform is by detecting a uniquely mapping novel peptide. However, the knowledge of the full-length protein isoforms expressed within a sample is not always possible when using short-read RNA-seq, which can return information on individual splice junctions but may not accurately define full-length transcripts.^17^ Long-read RNA-seq provides the full-length transcript and, by extension, the full- length protein isoform prediction; therefore, a novel peptide that directly maps to the full-length protein isoform lends support for its existence.

Using the sample-specific database, we discovered novel peptides for HUVECs, indicating that the reference proteome does not comprehensively capture all protein isoform diversity in a sample. We found 108 novel peptide sequences passing a global 1% FDR, for which they are not represented within the GENCODE or Uniprot databases (Figure 6A, Supplementary Information Table S6).^30,31^ Increased false positive rates for novel peptides have been observed previously^53^; therefore, we employed strict validation criteria for the novel peptides. Of the 108 novel peptides identified, 39 peptides had a Q-value score below 0.001, corresponding to a 0.1% global FDR. Upon manual annotation of these 39 peptides, we noted 30 peptides with especially strong spectral support, such as full ladders of b- and y- ion fragmentation peaks in the MS2 raw spectra. These novel peptides supported expression of novel alternative transcription or splicing events, such as retained intronic regions or novel exons (Figure 6B, Supplementary Information Table S6).

**Figure 6.**
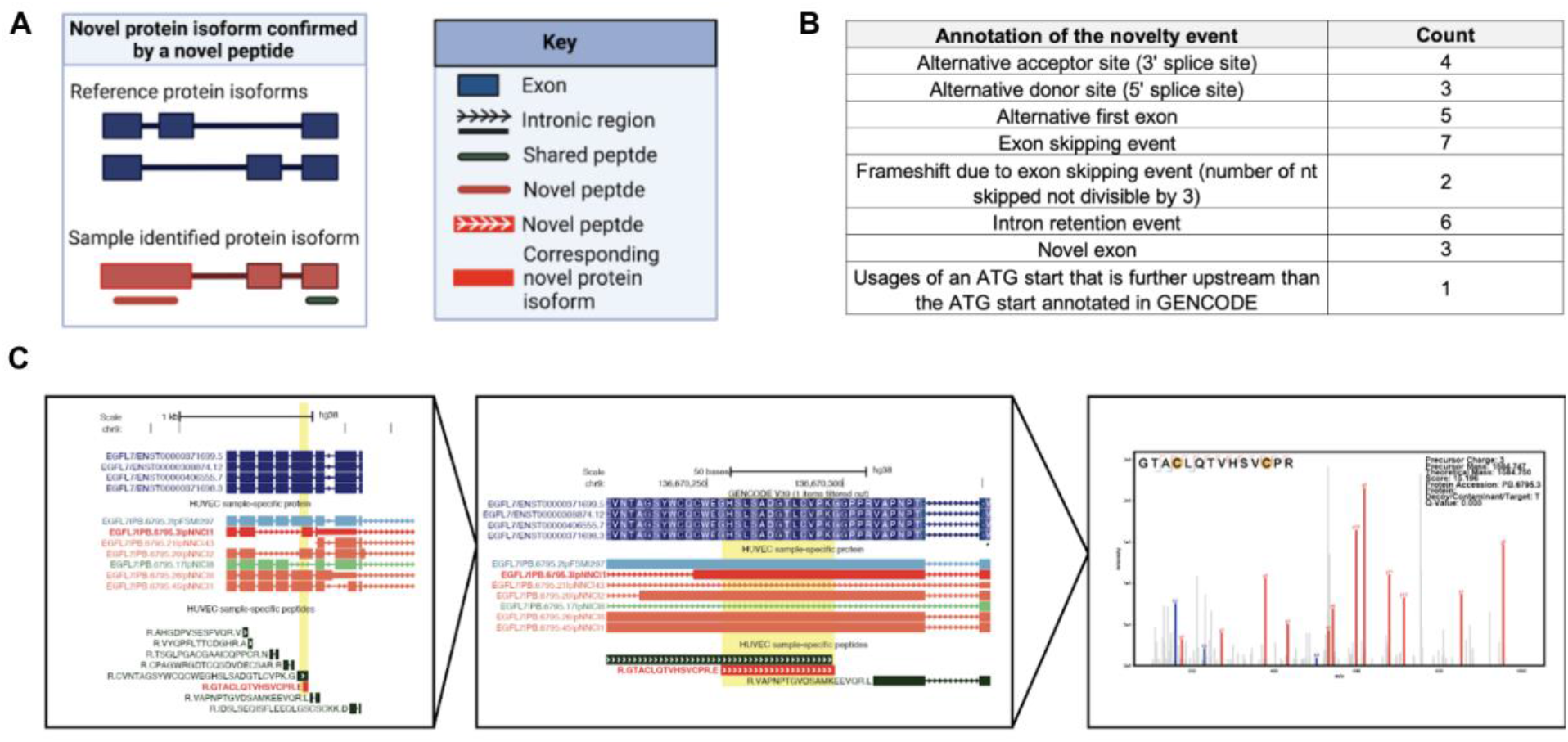
Novel protein isoforms discovered via unique peptides. (A) Novel protein isoform confirmed by identified novel peptides. (B) Table of the frequency of events supported confirmation of a novel peptide. (C) Novel peptide found for a protein isoform of endothelial gene *EGFL7*. Novel peptide and corresponding protein isoform shown in red, which supports a frameshift event for the protein isoform PB.6795.3.

Of the identified novel isoforms, we closely examined splicing events for genes previously implicated in endothelial pathways. Such novel isoforms could represent attractive candidate isoforms for further functional characterization. We found a novel peptide (Sequence: GTACLQTVHSVCPR) confirming the expression of a splice-induced frame-shifted region of *EGFL7* (Protein entry: PB.6795.3), a gene reported through the literature to be involved in vasculogenic pathways as well as hemogenic specification (Figure 6C).^54,55^ We also discovered two novel peptides for *PECAM1*. This is an important finding since *PECAM1* is a marker for endothelial cells and plays a role in the regulation of junctional integrity of endothelial cells and vascular barrier.^50^ Specifically, we discovered a novel peptide (Sequence: ELELLTSKDPPPSASQSAGITDLGKK, maps to protein entry PB.11293.45) corresponding to a novel exon, as well as a second novel peptide (Sequence: SDSGTYICTAEMLSQPR, mapping to protein entry PB.11293.25) that confirms the usage of a novel alternative donor site (Supporting information Table S6).

## Conclusions

Endothelial cells that line all blood vessels are critical for the cardiovascular system and their behaviors can be modulated by protein isoforms, though the extent of this mechanism is not known. To characterize isoform expression in endothelial cells, we performed long-read RNA sequencing (PacBio) of HUVECs to characterize transcript isoforms, and predicted proteins via their translation *in silico* to protein isoform sequences. To assess evidence for protein isoform expression, we performed MS analysis on the same HUVEC sample and used the HUVEC sample-specific database for MS searching. This general approach has been described and termed “long-read proteogenomics” to enhance protein isoform characterization.^22^

Our long-read proteogenomics workflow applied to HUVECs, led to the identification of 53,863 distinct transcript isoforms, of which 22,195 were novel. We also found 8,522 genes co-expressing multiple isoforms. Surprisingly, a quarter of the time, the most abundant isoform in HUVECs did not match the predicted “reference isoform” (GENCODE APPRIS principal isoform). This includes genes annotated in endothelial pathways including *CD34 and NRP1*. From the transcript sequences, we derived a hybrid protein isoform database that contains the highest confidence protein isoform predictions from PacBio-derived transcript isoform sequences, which was completed with GENCODE reference and contaminant sequences. The long-read-derived database captures almost all peptides and proteins detected from searches against the GENCODE protein database.

We identified 10,444 genes with peptide evidence. Based on the peptides identified through MS searching, we found support for expression of 4,451 genes based on uniquely mapping peptides. For the remaining 5,993 genes only evidenced by shared peptides, we incorporated the underlying transcript abundance information as an additional layer of evidence, nominating an additional 2,280 genes as potentially expressed. This group includes a novel isoform for endothelial gene *CDH5* (VE-Cadherin). This case exemplifies how a combination of the full-length transcript and proteomics data can lead to the discovery of novel protein isoforms that cannot be identified by MS data alone. We showed that the HUVEC sample-specific database enabled discovery of 108 novel protein isoforms based on novel peptide identifications. Among the novel protein isoforms identified is the endothelial gene *PECAM1*.

Our proteogenomic method shows promise for isoform discovery in endothelial cells, but opportunities exist for further improvements. First, limitations in the MS coverage mean that proteins with low abundance or poorly ionizable peptides remain undetected. Future work could involve targeted proteomics, such as parallel reaction monitoring or advanced targeted acquisition strategies, for sensitive detection of alternative protein isoforms.^56–58^ Second, the isoforms discovered in this study represent the results of a single cell line in a static culture condition. For the purposes of identifying isoforms that are dynamically regulated, multiple conditions should be examined. Third, the sample-specific database relies on the assumption that sequenced transcripts reflect protein sequences. Thus, we assume that transcripts are both fully sampled as well as moderately correlated to protein expression, which may not be the case for all genes.

Overall, we have shown the application of a long-read proteogenomics platform towards characterization of known and novel isoforms in primary endothelial cells. This approach can uncover isoform populations that could modulate endothelial cell phenotype and function. The systematic discovery of isoforms produces information to guide selection of candidate isoforms for functional studies. This approach can be extended to various endothelial cell contexts including both healthy and diseased states to chart isoforms changing across development or during onset of cardiovascular disease.

## Associated Content

### Supporting Information

Figure S1: Characterization of the HUVEC full-length transcriptome based on long-read RNA-seq data.

Table S1: Manually curated list of endothelial-relevant genes.

Table S2: Information on the major expressed isoform and APPRIS principle annotation for all and endothelial-related genes.

Figure S2: Derivation of predicted protein isoforms to generate a HUVEC sample-specific database.

Table S3: Databases derived from protein isoform models from the long-read proteogenomics. platform

Table S4: MetaMorpheus MS search parameters.

Figure S3. Basic pH HPLC fractions for the HUVEC MS data collection.

Figure S4: Comparison of proteomic coverage when using the HUVEC sample-specific versus UniProt protein databases for MS searching.

Table S5: Protein isoforms informed by shared and uniquely mapping peptide identifications.

Figure S5: Protein isoform informative peptides discovered when incorporating long-read data with all captured protein isoforms.

Table S6: Annotation of novel peptides detected from the sample-specific database.

### Author Contributions

GMS and MMM conceived of the project. GMS designed the study and supervised the project, along with KKH. MMM was involved in data collection, data analysis, interpretation, and conclusions, with discussions with GMS. LS performed the long-read RNA-seq experimental analysis. BTJ performed computational analysis, including long-read RNA-seq and proteomics analysis. MMM and EDJ performed the novel peptide analysis. JS performed computational analysis of novel peptides. BTJ, MMM and GMS contributed to analysis reproducibility, data curation and design of the workflow for the data described in this paper. MMM and GMS wrote the manuscript with input from all authors. All authors read and approved the final manuscript.

### Notes

The authors declare no competing financial interest.

## Acknowledgements

This work was financially supported by the Robert M. Berne Cardiovascular Research Center Training Program (T32HL007284) to MMM. We gratefully acknowledge the additional support by the NIH HL146056 and DK118728 to KKH. The long-read sequencing was performed at the Maryland Genomics at the University of Maryland Institute for Genome Science. Figures 1-6 were Created with Biorender.com. The graphical abstract was created with the help from Katharine Tuttle. We thank Dr. Gael Genet, Dr. Nicholas Chavkin, and Jordon Aragon for helpful comments and feedback related to this project.

## Abbreviations

AS: Alternative splicing
APPRIS: Annotation of principal and alternative splice isoforms
CCS: Circular consensus reads
CPM: Counts per million
GOrilla: Enriched GO terms
FDR: False-discovery-rate
FASTA: Fast-all
FASP: Filter-aided sample preparation
FSM: Full splice match
HUVEC: Human umbilical vein endothelial cells
PacBio processing: Iso-seq
MS: Mass-spectrometry
mRNA: Messenger RNA
NIC: Novel in catalog
NNC: Novel not in catalog
ORF: Open-reading frame
RNA-seq: RNA-sequencing
RNA-seq: Protein full splice match
pNIC: Protein novel in catalog
pNNC: Protein novel not in catalog
SMRT-seq: Single Molecule Real-Time sequencing
*SRSF5*: Serine and Arginine Rich Splicing Factor 5
SQANTI: Structural and Quality Annotation of Novel Transcript Isoforms
VEGF-A: Vascular endothelial growth factor A

## References

(1) Cleaver, O.; Melton, D. A. Endothelial Signaling during Development. Nat. Med. 2003, 9 (6), 661–668. https://doi.org/10.1038/nm0603-661.

(2) Rajendran, P.; Rengarajan, T.; Thangavel, J.; Nishigaki, Y.; Sakthisekaran, D.; Sethi, G.; Nishigaki, I. The Vascular Endothelium and Human Diseases. Int. J. Biol. Sci. 2013, 9 (10), 1057–1069. https://doi.org/10.7150/ijbs.7502.

(3) Richardson, M. R.; Lai, X.; Witzmann, F. A.; Yoder, M. C. Venous and Arterial Endothelial Proteomics: Mining for Markers and Mechanisms of Endothelial Diversity. Expert Rev. Proteomics 2010, 7 (6), 823–831. https://doi.org/10.1586/epr.10.92.

(4) Nordon, I.; Brar, R.; Hinchliffe, R.; Cockerill, G.; Loftus, I.; Thompson, M. The Role of Proteomic Research in Vascular Disease. J. Vasc. Surg. 2009, 49 (6), 1602–1612. https://doi.org/10.1016/j.jvs.2009.02.242.

(5) Farrokh, S.; Brillen, A. L.; Haendeler, J.; Altschmied, J.; Schaal, H. Critical Regulators of Endothelial Cell Functions: For a Change Being Alternative. Antioxid. Redox Signal. 2015, 22 (14), 1212–1229. https://doi.org/10.1089/ars.2014.6023.

(6) Bowler, E.; Oltean, S. Alternative Splicing in Angiogenesis. Int. J. Mol. Sci. 2019, 20 (9). https://doi.org/10.3390/ijms20092067.

(7) Mthembu, N. N.; Mbita, Z.; Hull, R.; Dlamini, Z. Abnormalities in Alternative Splicing of Angiogenesis-Related Genes and Their Role in HIV-Related Cancers. HIV AIDS 2017, 9, 77–93. https://doi.org/10.2147/HIV.S124911.

(8) Murphy, P. A.; Butty, V. L.; Boutz, P. L.; Begum, S.; Kimble, A. L.; Sharp, P. A.; Burge, C. B.; Hynes, R. O. Alternative RNA Splicing in the Endothelium Mediated in Part by Rbfox2 Regulates the Arterial Response to Low Flow. Elife 2018, 7. https://doi.org/10.7554/eLife.29494.

(9) Di Matteo, A.; Belloni, E.; Pradella, D.; Cappelletto, A.; Volf, N.; Zacchigna, S.; Ghigna, C. Alternative Splicing in Endothelial Cells: Novel Therapeutic Opportunities in Cancer Angiogenesis. J. Exp. Clin. Cancer Res. 2020, 39 (1), 275. https://doi.org/10.1186/s13046-020-01753-1.

(10) Hang, X.; Li, P.; Li, Z.; Qu, W.; Yu, Y.; Li, H.; Shen, Z.; Zheng, H.; Gao, Y.; Wu, Y.; Deng, M.; Sun, Z.; Zhang, C. Transcription and Splicing Regulation in Human Umbilical Vein Endothelial Cells under Hypoxic Stress Conditions by Exon Array. BMC Genomics 2009, 10, 126. https://doi.org/10.1186/1471-2164-10-126.

(11) Giampietro, C.; Deflorian, G.; Gallo, S.; Di Matteo, A.; Pradella, D.; Bonomi, S.; Belloni, E.; Nyqvist, D.; Quaranta, V.; Confalonieri, S.; Bertalot, G.; Orsenigo, F.; Pisati, F.; Ferrero, E.; Biamonti, G.; Fredrickx, E.; Taveggia, C.; Wyatt, C. D.; Irimia, M.; Di Fiore, P. P.; Blencowe, J.; Dejana, E.; Ghigna, C. The Alternative Splicing Factor Nova2 Regulates Vascular Development and Lumen Formation. Nat. Commun. 2015, 6, 8479. https://doi.org/10.1038/ncomms9479.

(12) Khan, S.; Taverna, F.; Rohlenova, K.; Treps, L.; Geldhof, V.; de Rooij, L.; Sokol, L.; Pircher, A.; Conradi, L.-C.; Kalucka, J.; Schoonjans, L.; Eelen, G.; Dewerchin, M.; Karakach, T.; Li, X.; Goveia, J.; Carmeliet, P. EndoDB: A Database of Endothelial Cell Transcriptomics Data. Nucleic Acids Res. 2019, 47 (D1), D736–D744. https://doi.org/10.1093/nar/gky997.

(13) Mudge, J. M.; Harrow, J. The State of Play in Higher Eukaryote Gene Annotation. Nat. Rev. Genet. 2016, 17 (12), 758–772. https://doi.org/10.1038/nrg.2016.119.

(14) Caniuguir, A.; Krause, B. J.; Hernandez, C.; Uauy, R.; Casanello, P. Markers of Early Endothelial Dysfunction in Intrauterine Growth Restriction-Derived Human Umbilical Vein Endothelial Cells Revealed by 2D-DIGE and Mass Spectrometry Analyses. Placenta 2016, 41, 14–26. https://doi.org/10.1016/j.placenta.2016.02.016.

(15) Banarjee, R.; Sharma, A.; Bai, S.; Deshmukh, A.; Kulkarni, M. Proteomic Study of Endothelial Dysfunction Induced by AGEs and Its Possible Role in Diabetic Cardiovascular Complications. J. Proteomics 2018, 187, 69–79. https://doi.org/10.1016/j.jprot.2018.06.009.

(16) Madugundu, A. K.; Na, C. H.; Nirujogi, R. S.; Renuse, S.; Kim, K. P.; Burns, K. H.; Wilks, C.; Langmead, B.; Ellis, S. E.; Collado-Torres, L.; Halushka, M. K.; Kim, M. S.; Pandey, A. Integrated Transcriptomic and Proteomic Analysis of Primary Human Umbilical Vein Endothelial Cells. Proteomics 2019, 19 (15), e1800315. https://doi.org/10.1002/pmic.201800315.

(17) Steijger, T.; Abril, J. F.; Engstrom, P. G.; Kokocinski, F.; Rgasp Consortium; Hubbard, T. J., Guigo, R.; Harrow, J.; Bertone, P. Assessment of Transcript Reconstruction Methods for RNA-Seq. Nat. Methods 2013, 10 (12), 1177–1184. https://doi.org/10.1038/nmeth.2714.

(18) Nesvizhskii, A. I. Proteogenomics: Concepts, Applications and Computational Strategies. Nat. Methods 2014, 11 (11), 1114–1125. https://doi.org/10.1038/nmeth.3144.

(19) van Dijk, E. L.; Jaszczyszyn, Y.; Naquin, D.; Thermes, C. The Third Revolution in Sequencing Technology. Trends Genet. 2018, 34 (9), 666–681. https://doi.org/10.1016/j.tig.2018.05.008.

(20) Eid, J.; Fehr, A.; Gray, J.; Luong, K.; Lyle, J.; Otto, G.; Peluso, P.; Rank, D.; Baybayan, P.; Bettman, B.; Bibillo, A.; Bjornson, K.; Chaudhuri, B.; Christians, F.; Cicero, R.; Clark, S.; Dalal, R.; Dewinter, A.; Dixon, J.; Foquet, M.; Gaertner, A.; Hardenbol, P.; Heiner, C.; Hester, K.; Holden, D.; Kearns, G.; Kong, X.; Kuse, R.; Lacroix, Y.; Lin, S.; Lundquist, P.; Ma, C.; Marks, P.; Maxham, M.; Murphy, D.; Park, I.; Pham, T.; Phillips, M.; Roy, J.; Sebra, R.; Shen, G.; Sorenson, J.; Tomaney, A.; Travers, K.; Trulson, M.; Vieceli, J.; Wegener, J.; Wu, D.; Yang, A.; Zaccarin, D.; Zhao, P.; Zhong, F.; Korlach, J.; Turner, S. Real-Time DNA Sequencing from Single Polymerase Molecules. Science 2009, 323 (5910), 133–138. https://doi.org/10.1126/science.1162986.

(21) Rhoads, A.; Au, K. F. PacBio Sequencing and Its Applications. Genomics Proteomics Bioinformatics 2015, 13 (5), 278–289. https://doi.org/10.1016/j.gpb.2015.08.002.

(22) Miller, R. M.; Jordan, B. T.; Mehlferber, M. M.; Jeffery, E. D.; Chatzipantsiou, C.; Kaur, S.; Millikin, R. J.; Dai, Y.; Tiberi, S.; Castaldi, P. J.; Shortreed, M. R.; Luckey, C. J.; Conesa, A.; Smith, L. M.; Deslattes Mays, A.; Sheynkman, G. M. Enhanced Protein Isoform Characterization through Long-Read Proteogenomics. Genome Biol. 2022, 23 (1), 1–28. https://doi.org/10.1186/s13059-022-02624-y.

(23) Sharon, D.; Tilgner, H.; Grubert, F.; Snyder, M. A Single-Molecule Long-Read Survey of the Human Transcriptome. Nat. Biotechnol. 2013, 31 (11), 1009–1014. https://doi.org/10.1038/nbt.2705.

(24) Deslattes Mays, A.; Schmidt, M.; Graham, G.; Tseng, E.; Baybayan, P.; Sebra, R.; Sanda, M.; Mazarati, J. B.; Riegel, A.; Wellstein, A. Single-Molecule Real-Time (SMRT) Full-Length RNA-Sequencing Reveals Novel and Distinct mRNA Isoforms in Human Bone Marrow Cell Subpopulations. Genes 2019, 10 (4), 17. https://doi.org/10.3390/genes10040253.

(25) Verbruggen, S.; Gessulat, S.; Gabriels, R.; Matsaroki, A.; Van de Voorde, H.; Kuster, B.; Degroeve, S.; Martens, L.; Van Criekinge, W.; Wilhelm, M.; Menschaert, G. Spectral Prediction Features as a Solution for the Search Space Size Problem in Proteogenomics. Mol. Cell. Proteomics 2021, 2021/04/07, 100076. https://doi.org/10.1016/j.mcpro.2021.100076.

(26) Anvar, S. Y.; Allard, G.; Tseng, E.; Sheynkman, G. M.; de Klerk, E.; Vermaat, M.; Yin, R. H.; Johansson, H. E.; Ariyurek, Y.; den Dunnen, J. T.; Turner, S. W.; ‘t Hoen, P. A. Full-Length mRNA Sequencing Uncovers a Widespread Coupling between Transcription Initiation and mRNA Processing. Genome Biology. 2018. https://doi.org/10.1186/s13059-018-1418-0.

(27) Nesvizhskii, A. I.; Aebersold, R. Interpretation of Shotgun Proteomic Data. Mol. Cell. Proteomics 2005, 4 (10), 1419–1440. https://doi.org/10.1074/mcp.R500012-MCP200.

(28) Gordon, S. P.; Tseng, E.; Salamov, A.; Zhang, J.; Meng, X.; Zhao, Z.; Kang, D.; Underwood, J.; Grigoriev, I. V.; Figueroa, M.; Schilling, J. S.; Chen, F.; Wang, Z. Widespread Polycistronic Transcripts in Fungi Revealed by Single-Molecule mRNA Sequencing. PLoS One 2015, 10 (7). https://doi.org/10.1371/journal.pone.0132628.

(29) Tardaguila, M.; de la Fuente, L., Marti, C.; Pereira, C.; Pardo-Palacios, F. J.; Del Risco, H.; Ferrell, M.; Mellado, M.; Macchietto, M.; Verheggen, K.; Edelmann, M.; Ezkurdia, I.; Vazquez, J.; Tress, M.; Mortazavi, A.; Martens, L.; Rodriguez-Navarro, S.; Moreno-Manzano, V.; Conesa, A. SQANTI: Extensive Characterization of Long-Read Transcript Sequences for Quality Control in Full-Length Transcriptome Identification and Quantification. Genome Res. 2018, 2018/02/15. https://doi.org/10.1101/gr.222976.117.

(30) Frankish, A.; Diekhans, M.; Ferreira, A. M.; Johnson, R.; Jungreis, I.; Loveland, J.; Mudge, J. M.; Sisu, C.; Wright, J.; Armstrong, J.; Barnes, I.; Berry, A.; Bignell, A.; Sala, S. C.; Chrast, J.; Cunningham, F.; Di Domenico, T.; Donaldson, S.; Fiddes, I. T.; Giron, C. G.; Gonzalez, J. M.; Grego, T.; Hardy, M.; Hourlier, T.; Hunt, T.; Izuogu, O. G.; Lagarde, J.; Martin, F. J.; Martinez, L.; Mohanan, S.; Muir, P.; Navarro, F. C. P.; Parker, A.; Pei, B. K.; Pozo, F.; Ruffier, M.; Schmitt, B. M.; Stapleton, E.; Suner, M. M.; Sycheva, I.; Uszczynska-Ratajczak, B., Xu, J.; Yates, A.; Zerbino, D.; Zhang, Y.; Aken, B.; Choudhary, J. S.; Gerstein, M.; Guigo, R.; Hubbard, T. J.; Kellis, M.; Paten, B.; Reymond, A.; Tress, M. L.; Flicek, P. GENCODE Reference Annotation for the Human and Mouse Genomes. Nucleic Acids Res. 2019, 47 (D1), D766–D773. https://doi.org/10.1093/nar/gky955.

(31) UniProt, Consortium. UniProt: A Worldwide Hub of Protein Knowledge. Nucleic Acids Res. 2019, 47 (D1), D506–D515. https://doi.org/10.1093/nar/gky1049.

(32) McGarvey, P. B.; Nightingale, A.; Luo, J.; Huang, H.; Martin, M. J.; Wu, C.; UniProt, Consortium. UniProt Genomic Mapping for Deciphering Functional Effects of Missense Variants. Hum. Mutat. 2019, 40 (6), 694–705. https://doi.org/10.1002/humu.23738.

(33) Huang, K. K.; Huang, J.; Wu, J. K. L.; Lee, M.; Tay, S. T.; Kumar, V.; Ramnarayanan, K.; Padmanabhan, N.; Xu, C.; Tan, A. L. K.; Chan, C.; Kappei, D.; Göke, J.; Tan, P. Long-Read Transcriptome Sequencing Reveals Abundant Promoter Diversity in Distinct Molecular Subtypes of Gastric Cancer. Genome Biol. 2021, 22 (1), 44. https://doi.org/10.1186/s13059-021-02261-x.

(34) Leung, S. K.; Jeffries, A. R.; Castanho, I.; Jordan, B. T.; Moore, K.; Davies, J. P.; Dempster, E. L.; Bray, N. J.; O’Neill, P.; Tseng, E.; Ahmed, Z.; Collier, D. A.; Jeffery, E. D.; Prabhakar, S.; Schalkwyk, L.; Jops, C.; Gandal, M. J.; Sheynkman, G. M.; Hannon, E.; Mill, J. Full-Length Transcript Sequencing of Human and Mouse Cerebral Cortex Identifies Widespread Isoform Diversity and Alternative Splicing. Cell Rep. 2021, 37 (7), 110022. https://doi.org/10.1016/j.celrep.2021.110022.

(35) Marcelo, K. L.; Goldie, L. C.; Hirschi, K. K. Regulation of Endothelial Cell Differentiation and Specification. Circ. Res. 2013, 112 (9), 1272–1287. https://doi.org/10.1161/CIRCRESAHA.113.300506.

(36) Aragon, J. W.; Hirschi, K. K. Endothelial Cell Differentiation and Hemogenic Specification. Cold Spring Harb. Perspect. Med. 2022. https://doi.org/10.1101/cshperspect.a041164.

(37) Eden, E.; Navon, R.; Steinfeld, I.; Lipson, D.; Yakhini, Z. GOrilla: A Tool for Discovery and Visualization of Enriched GO Terms in Ranked Gene Lists. BMC Bioinformatics 2009, 10, 48. https://doi.org/10.1186/1471-2105-10-48.

(38) Kofler, N. M.; Simons, M. Angiogenesis versus Arteriogenesis: Neuropilin 1 Modulation of VEGF Signaling. F1000Prime Rep. 2015, 7, 26. https://doi.org/10.12703/P7-26.

(39) Lanahan, A.; Zhang, X.; Fantin, A.; Zhuang, Z.; Rivera-Molina, F.; Speichinger, K.; Prahst, C.; Zhang, J.; Wang, Y.; Davis, G.; Toomre, D.; Ruhrberg, C.; Simons, M. The Neuropilin 1 Cytoplasmic Domain Is Required for VEGF-A-Dependent Arteriogenesis. Dev. Cell 2013, 25 (2), 156–168. https://doi.org/10.1016/j.devcel.2013.03.019.

(40) Rodriguez, J. M.; Rodriguez-Rivas, J.; Di Domenico, T.; Vázquez, J.; Valencia, A.; Tress, M. L. APPRIS 2017: Principal Isoforms for Multiple Gene Sets. Nucleic Acids Res. 2018, 46 (D1), D213–D217. https://doi.org/10.1093/nar/gkx997.

(41) Wang, X.; Slebos, R. J.; Wang, D.; Halvey, P. J.; Tabb, D. L.; Liebler, D. C.; Zhang, B. Protein Identification Using Customized Protein Sequence Databases Derived from RNA-Seq Data. J. Proteome Res. 2012, 11 (2), 1009–1017. https://doi.org/10.1021/pr200766z.

(42) Elias, J. E.; Gygi, S. P. Target-Decoy Search Strategy for Increased Confidence in Large-Scale Protein Identifications by Mass Spectrometry. Nat. Methods 2007, 4 (3), 207–214. https://doi.org/10.1038/nmeth1019.

(43) Solntsev, S. K.; Shortreed, M. R.; Frey, B. L.; Smith, L. M. Enhanced Global Post-Translational Modification Discovery with MetaMorpheus. J. Proteome Res. 2018, 17 (5), 1844–1851. https://doi.org/10.1021/acs.jproteome.7b00873.

(44) Wang, D.; Eraslan, B.; Wieland, T.; Hallstrom, B.; Hopf, T.; Zolg, D. P.; Zecha, J.; Asplund, A.; Li, L. H.; Meng, C.; Frejno, M.; Schmidt, T.; Schnatbaum, K.; Wilhelm, M.; Ponten, F.; Uhlen, M.; Gagneur, J.; Hahne, H.; Kuster, B. A Deep Proteome and Transcriptome Abundance Atlas of 29 Healthy Human Tissues. Mol. Syst. Biol. 2019, 15 (2), e8503. https://doi.org/10.15252/msb.20188503.

(45) Blakeley, P.; Siepen, J. A.; Lawless, C.; Hubbard, S. J. Investigating Protein Isoforms via Proteomics: A Feasibility Study. Proteomics 2010, 10 (6), 1127–1140. https://doi.org/10.1002/pmic.200900445.

(46) Lau, E.; Han, Y.; Williams, D. R.; Thomas, C. T.; Shrestha, R.; Wu, J. C.; Lam, M. P. Y. Splice-Junction-Based Mapping of Alternative Isoforms in the Human Proteome. Cell Rep. 2019, 29 (11), 3751–3765.e5. https://doi.org/10.1016/j.celrep.2019.11.026.

(47) Carlyle, B. C.; Kitchen, R. R.; Zhang, J.; Wilson, R. S.; Lam, T. T.; Rozowsky, J. S.; Williams, K. R.; Sestan, N.; Gerstein, M. B.; Nairn, A. C. Isoform-Level Interpretation of High-Throughput Proteomics Data Enabled by Deep Integration with RNA-Seq. J. Proteome Res. 2018, 17 (10), 3431–3444. https://doi.org/10.1021/acs.jproteome.8b00310.

(48) Salovska, B.; Zhu, H.; Gandhi, T.; Frank, M.; Li, W.; Rosenberger, G.; Wu, C.; Germain, P.- L.; Zhou, H.; Hodny, Z.; Reiter, L.; Liu, Y. Isoform-Resolved Correlation Analysis between mRNA Abundance Regulation and Protein Level Degradation. Mol. Syst. Biol. 2020, 16 (3), e9170. https://doi.org/10.15252/msb.20199170.

(49) Sauteur, L.; Krudewig, A.; Herwig, L.; Ehrenfeuchter, N.; Lenard, A.; Affolter, M.; Belting, H.-G. Cdh5/VE-Cadherin Promotes Endothelial Cell Interface Elongation via Cortical Actin Polymerization during Angiogenic Sprouting. Cell Rep. 2014, 9 (2), 504–513. https://doi.org/10.1016/j.celrep.2014.09.024.

(50) Privratsky, J. R.; Newman, P. J. PECAM-1: Regulator of Endothelial Junctional Integrity. Cell Tissue Res. 2014, 355 (3), 607–619. https://doi.org/10.1007/s00441-013-1779-3.

(51) Cao, G.; O’Brien, C. D.; Zhou, Z.; Sanders, S. M.; Greenbaum, J. N.; Makrigiannakis, A.; DeLisser, H. M. Involvement of Human PECAM-1 in Angiogenesis and in Vitro Endothelial Cell Migration. Am. J. Physiol. Cell Physiol. 2002, 282 (5), C1181–C1190. https://doi.org/10.1152/ajpcell.00524.2001.

(52) Dusserre, N.; L’Heureux, N.; Bell, K. S.; Stevens, H. Y.; Yeh, J.; Otte, L. A.; Loufrani, L.; Frangos, J. A. PECAM-1 Interacts with Nitric Oxide Synthase in Human Endothelial Cells: Implication for Flow-Induced Nitric Oxide Synthase Activation. Arterioscler. Thromb. Vasc. Biol. 2004, 24 (10), 1796–1802. https://doi.org/10.1161/01.ATV.0000141133.32496.41.

(53) Castellana, N.; Bafna, V. Proteogenomics to Discover the Full Coding Content of Genomes: A Computational Perspective. J. Proteomics 2010, 73 (11), 2124–2135. https://doi.org/10.1016/j.jprot.2010.06.007.

(54) Nichol, D.; Stuhlmann, H. EGFL7: A Unique Angiogenic Signaling Factor in Vascular Development and Disease. Blood 2012, 119 (6), 1345–1352. https://doi.org/10.1182/blood-2011-10-322446.

(55) Schmidt, M. H. H.; Bicker, F.; Nikolic, I.; Meister, J.; Babuke, T.; Picuric, S.; Müller-Esterl, W.; Plate, K. H.; Dikic, I. Epidermal Growth Factor-like Domain 7 (EGFL7) Modulates Notch Signalling and Affects Neural Stem Cell Renewal. Nat. Cell Biol. 2009, 11 (7), 873–880. https://doi.org/10.1038/ncb1896.

(56) Gallien, S.; Kim, S. Y.; Domon, B. Large-Scale Targeted Proteomics Using Internal Standard Triggered-Parallel Reaction Monitoring (IS-PRM)*[S]. Mol. Cell. Proteomics 2015, 14 (6), 1630–1644.

(57) Erickson, B. K.; Rose, C. M.; Braun, C. R.; Erickson, A. R.; Knott, J.; McAlister, G. C.; Wühr, M.; Paulo, J. A.; Everley, R. A.; Gygi, S. P. A Strategy to Combine Sample Multiplexing with Targeted Proteomics Assays for High-Throughput Protein Signature Characterization. Mol. Cell 2017, 65 (2), 361–370. https://doi.org/10.1016/j.molcel.2016.12.005.

(58) Wichmann, C.; Meier, F.; Winter, S. V.; Brunner, A.-D.; Cox, J.; Mann, M. MaxQuant.Live Enables Global Targeting of More than 25,000 Peptides. https://doi.org/10.1101/443838.

(59) Wiśniewski, J. R. Filter-Aided Sample Preparation for Proteome Analysis. Methods in Molecular Biology. 2018, pp 3–10. https://doi.org/10.1007/978-1-4939-8695-8_1.

(60) Wang, L.; Park, H. J.; Dasari, S.; Wang, S.; Kocher, J. P.; Li, W. CPAT: Coding-Potential Assessment Tool Using an Alignment-Free Logistic Regression Model. Nucleic Acids Res. 2013, 41 (6), e74. https://doi.org/10.1093/nar/gkt006.

(61) Smith, L. M.; Agar, J. N.; Chamot-Rooke, J.; Danis, P. O.; Ge, Y.; Loo, J. A.; Paša-Tolić, L.; Tsybin, Y. O.; Kelleher, N. L.; Consortium for Top-Down Proteomics. The Human Proteoform Project: Defining the Human Proteome. Sci Adv 2021, 7 (46), eabk0734. https://doi.org/10.1126/sciadv.abk0734.

