## Supporting Information for "Characterization of protein isoform diversity in human umbilical vein endothelial cells (HUVECs) via long-read proteogenomics"

### Table of Contents:

S-1: Cover sheet

S-2: Supplementary Figure S1. Characterization of the HUVEC full-length transcriptome based on long-read RNA-seq data.

S-3: Supplementary Table S1. Manually curated list of endothelial-relevant genes

S-4: Supplementary Table S2. Information on the major expressed isoform and APPRIS principal annotation for all and endothelial-related genes

S-5: Supplementary Figure S2. Derivation of predicted protein isoforms to generate a HUVEC sample-specific database.

S-6: Supplementary Table S3. Databases derived from protein isoform models from the long-read proteogenomics platform

S-7: Supplementary Table S4. Metaorpheus MS search parameters

S-8: Supplementary Figure S3. Basic pH HPLC fractions for the HUVEC MS data collection.

S-9: Supplementary Figure S4. Comparison of proteomic coverage when using the HUVEC sample-specific versus UniProt protein databases for MS searching.

S-10 Supplementary Figure S5. Plectin (PLEC) gene evidenced by seven unique isoforms.

S-11: Supplementary Table S5: Protein isoforms informed by shared and uniquely mapping peptide identifications.

S-12: Supplementary Table S6: Annotation of novel peptides detected from the sample-specific database

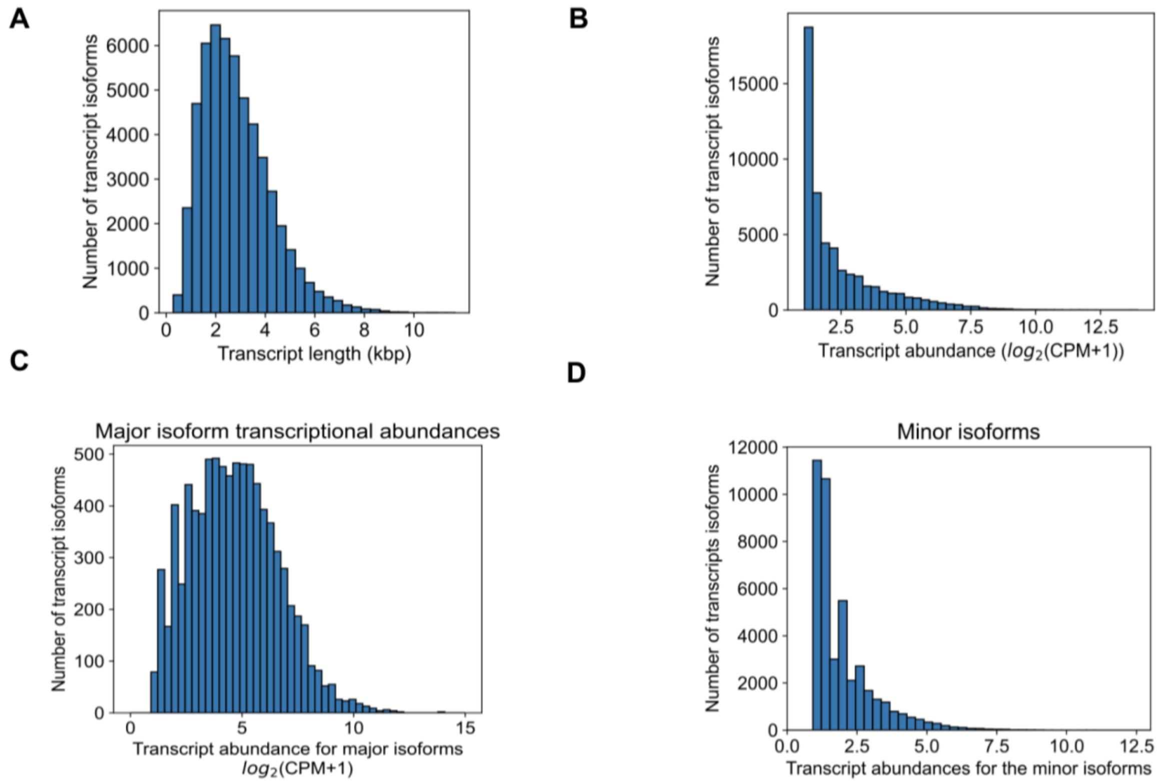


**Figure S1.** Characterization of the HUVEC full-length transcriptome based on long-read RNA-seq data. A**)** Distribution of transcript isoform length B) Distribution of the transcript abundance (counts per million, CPM) B) Distribution of all transcripts identified from the Iso-Seq pipeline with abundance greater than one CPM. C) Distribution of the abundances foro the most highly expressed isoform for each gene, i.e., the major isoform. D) Distribution of the abundances of the minor isoforms reported for all transcripts with CPM>1.

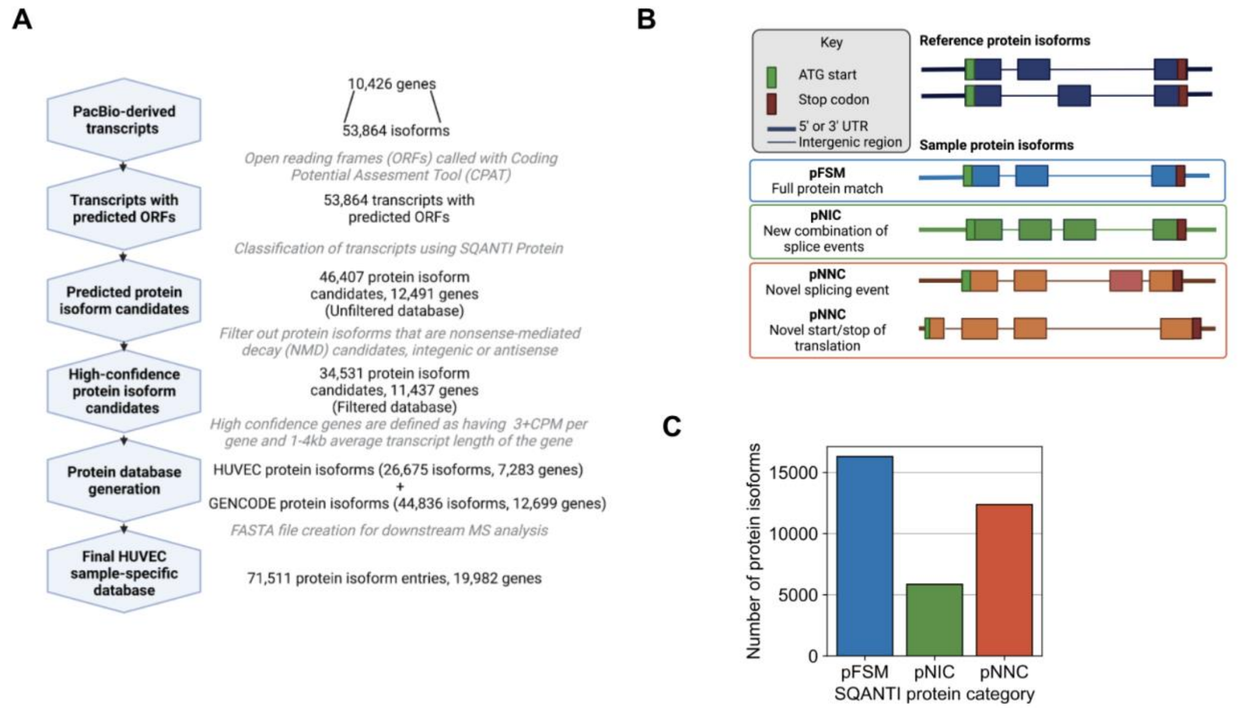


**Figure S2.** Derivation of predicted protein isoforms to generate a HUVEC sample-specific database.. (A) Schematic of protein database generation from long-read RNA-seq data. (B) Schematic of SQANTI Protein classification (C) Bar chart indicating the frequency of different protein isoforms based on novelty category.

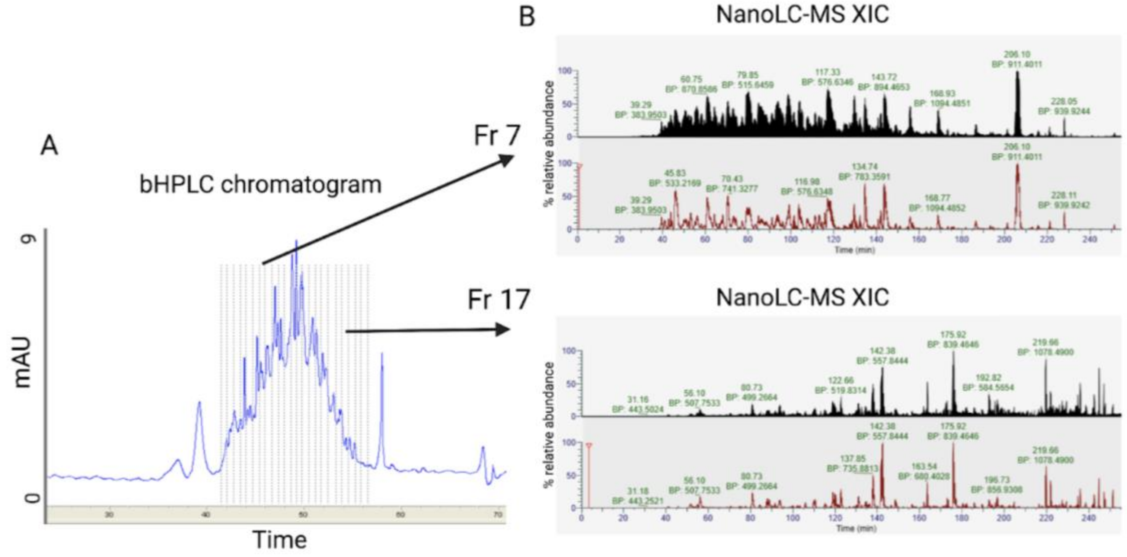


**Figure S3.** Basic pH HPLC fractions for the HUVEC MS data collection. . (A) UV trace of peptide elution from offline fractionation (range of fractions shown with dashed lines). (B) Total ion (black) and base peak (red) chromatograms from LC-MS analysis of peptides from representative fractions 7 and 17. Basic High Performance Liquid Chromatography (bHPLC), Extracted ion chromatogram (XIC), Nanoscale liquid chromatography coupled to tandem mass spectrometry (nano LC–MS/MS), milli-Absorbance Units (mAU).

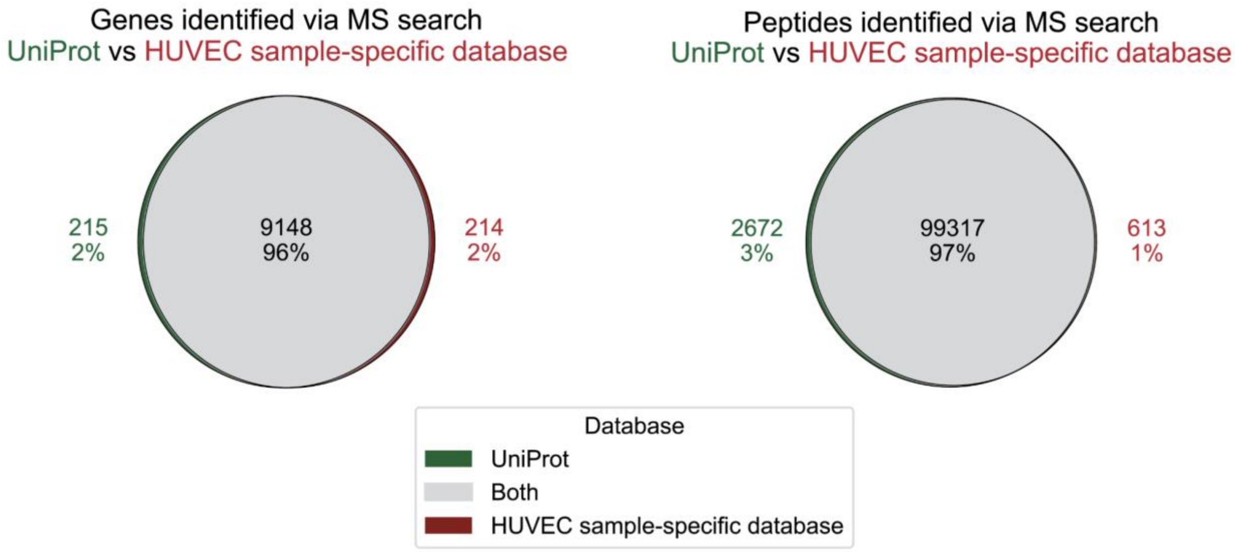


**Figure S4.** Comparison of proteomic coverage when using the HUVEC sample-specific versus UniProt protein databases for MS searching.

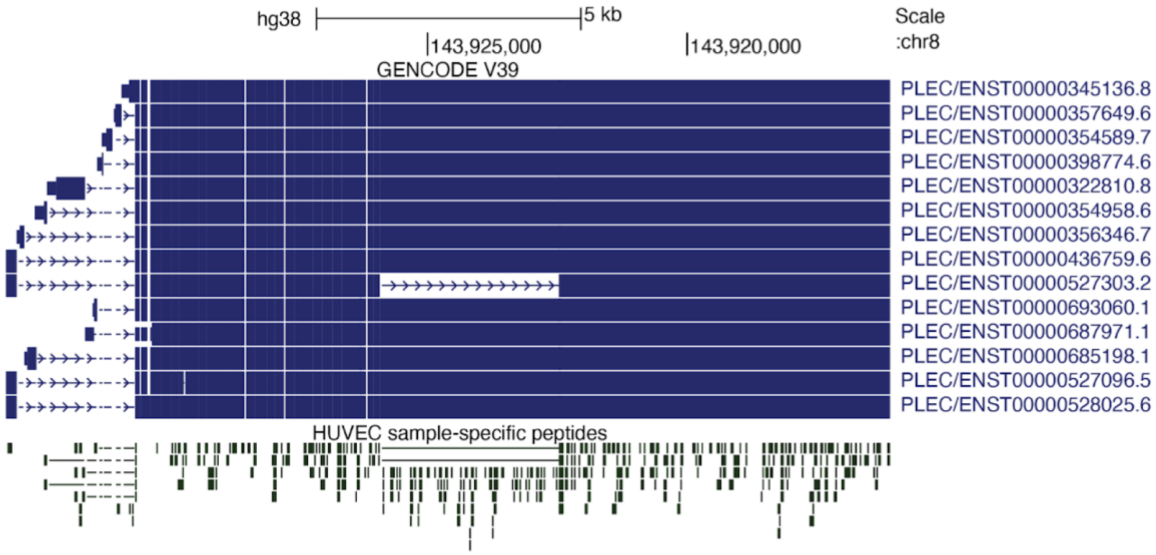


**Figure S5.** Plectin (*PLEC*) gene evidenced by seven unique isoforms. Plectin (*PLEC*) gene in which MS analysis identified seven unique isoforms each evidenced by their own unique peptide.
